# Experience-Driven Rate Modulation is Reinstated During Hippocampal Replay

**DOI:** 10.1101/2021.07.15.452506

**Authors:** M. Tirole, M. Huelin Gorriz, M. Takigawa, L. Kukovska, D. Bendor

## Abstract

Replay, the sequential reactivation within a neuronal ensemble, is a central hippocampal mechanism postulated to drive memory processing. While both rate and place representations are used by hippocampal place cells to encode behavioral episodes, replay has been largely defined by only the latter – based on the fidelity of sequential activity across neighboring place fields. Here we show that dorsal CA1 place cells in rats can modulate their firing rate between replay events of two different contexts. This experience-dependent phenomenon mirrors the same pattern of rate modulation observed during behavior and can be used independently from place information within replay sequences to discriminate between contexts. Our results reveal the existence of two complementary neural representations available for memory processes.

## INTRODUCTION

A key function of the hippocampus is the initial encoding and subsequent consolidation of episodic memories. Hippocampal place cells are activated in sequential patterns during behavioral episodes (e.g. a rodent running towards a goal), with each neuron’s activity modulated by the animal’s position within an environment (i.e. a place field) (O’Keefe & Dostrovsky, 1971). During offline periods such as sleep, place cells spontaneously reactivate, replaying the same sequential pattern previously activated during the behavioral episode (Lee & Wilson, 2002; Wilson & McNaughton, 1994). This phenomenon of neural replay is postulated to drive systems-level memory consolidation (Crowley, Bendor, & Javadi, 2019; Fernández-Ruiz et al., 2019; Girardeau, Benchenane, Wiener, Buzsáki, & Zugaro, 2009; Gridchyn, Schoenenberger, ONeill, & Csicsvari, 2020). Hippocampal replay has largely been defined by the sequential reactivation of place cell ensembles (Davidson, Kloosterman, & Wilson, 2009; Foster & Wilson, 2006; Genzel et al., 2020; Lee & Wilson, 2002), driven by both where and when each place field is activated during a behavioral episode (here referred to as place representation). Yet, place cells also carry additional information in the magnitude of their place field’s activity (here referred to as rate representation). The magnitude of a place field’s activity (i.e. peak in-field firing rate) is modulated by both local and global contextual cues (Leutgeb et al., 2005; Ravassard et al., 2013; Zhao, Wang, Spruston, & Magee, 2020), as well as behavioral variables (e.g. animal’s speed during locomotion) (Huxter, Burgess, & O’Keefe, 2003), and cognitive events (e.g. animal’s attention and perception) (Monaco, Rao, Roth, & Knierim, 2014; Olypher, Lánský, & Fenton, 2002). However, we do not know whether the hippocampus is capable of reinstating place cell rate modulation during replay. How hippocampal replay exactly contributes to consolidating episodic memories remains an open question. Examining whether rate modulation is reinstated during replay will help discern whether replay only represents possible transitions between adjacent locations: that is, the order of place fields within a spatial trajectory. Alternatively, replay could serve a larger role, with the capacity to reinstate a near replica of neural activity previously driven during behavior, providing a richer, more complete memory trace.

## RESULTS

### Place cells reinstate their rate modulation during replay

To address this question, we recorded extracellularly from the dorsal CA1 of rats (n= 5, male Lister-hooded), trained to run back and forth along two novel linear tracks (2m) to collect a liquid reward at each end (Figure 1A, Figure S1A, Table S1A, and see Methods). A view-obstructing divider was placed between tracks and distinguishing visual cues were present within each environment to facilitate contextual discrimination and, in turn, hippocampal remapping. Additionally, rats rested in a quiet remote location, both before (PRE) and after (POST) the exploration of the two tracks.

**Figure 1.**
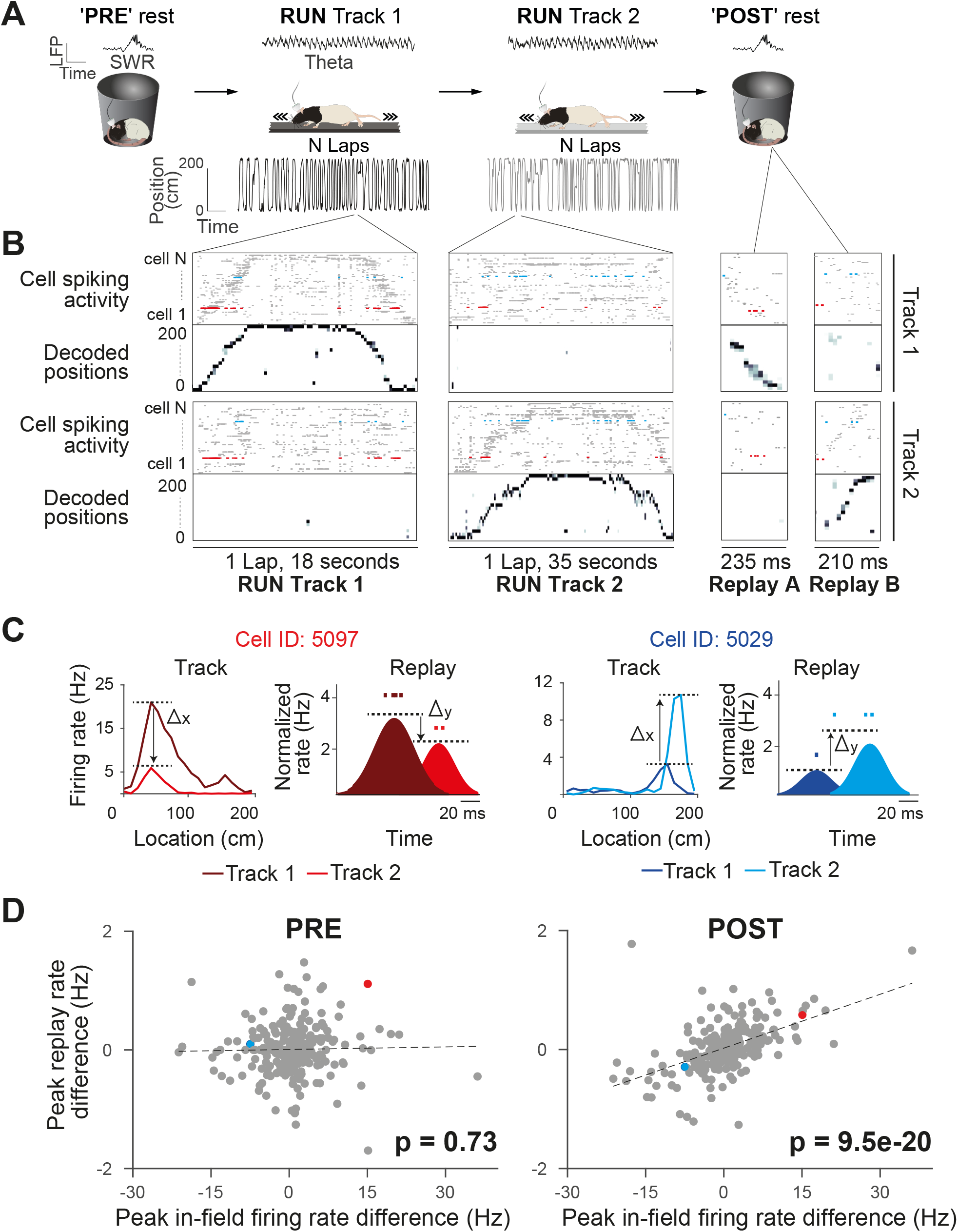
Rate coding in wake and rest. (A) Experimental design. Each recording session, the rats rested (PRE), ran back and forth on two different linear tracks (RUN), and then rested again (POST) (left to right). An example of LFP activity in each behavioral state is displayed [RUN-theta, PRE/POST-sharp wave ripples (SWR)]. (B) Raster plots of spiking activity of cells sorted by their peak firing rate on the track (top: Track 2, n = 57 cells; bottom: Track 1, n = 72 cells), and the associated decoded positions for a single exemplar lap (first two columns) and replay event (last two columns), for each track (top: Track 2; bottom: Track 1). Example from Rat 3, session 2. (C) Two example neurons (highlighted in B) that changed their peak instantaneous firing rate (shaded areas, cartoon depictions of instantaneous firing rates) during two example replay events (displayed in B) in the same direction as their peak in-field firing rate difference between tracks (unshaded areas, actual ratemaps). The activity and properties of the example neurons are color coded throughout all the panels. (D) The peak in-field firing rate difference (Track1-Track2) significantly predicts the average peak instantaneous replay rate difference (Track1-Track2) in POST rest, but not PRE rest. Each data point is a neuron active on both tracks (PRE: n = 254cells, POST: n = 275cells).

We observed global remapping of place cells between the two linear tracks (Pearson correlation coefficient (r) between tracks population vectors, r = 0.11 ± 0.13), such that the majority of place fields shifted their location along the track and/or modulated their firing rate (i.e. changed to higher or lower peak in-field firing rate) (Figure S1B). Both the context (which linear track) and current position (location of the animal on the linear track) could be inferred from place cell activity with high accuracy using a naïve Bayes decoder (Figure 1B and Figure S1C). Track detection accuracy exceeded 80% during RUN periods on both tracks (all sessions: 91 ± 5 %) with an average median error of 5cm (all sessions: 6.08 ± 2.5 cm). A population vector analysis showed high correlation between the ratemaps of the first and second half of running within each track (Pearson correlation coefficient (r) within track population vectors, r= 0.81 ± 0.11), suggesting high place field stability over run. We also used Bayesian decoding to detect replay sequences of each linear track (Figure 1B), with statistically significant replay trajectories identified by comparing the weighted correlation score obtained from the posterior probabilities of each decoded event to three types of event shuffles (Figure S6; Total replay events = 5800, PRE: Track1= 51±33, Track2= 46±22, POST: Track1= 151±52, Track2= 159±58, RUN: Track1= 67±20, Track2= 78±24; see Methods).

First, we sought to investigate whether the contextually-driven place fields’ rate modulation between tracks is conserved during offline replay. We use the term “contextually-driven” to indicate a rate change occurring between the track 1 and track 2 contexts, albeit driven by environmental cues, track position, and/or behavioral differences in the rat. Using all place cells active on both tracks and stable across the whole run (Figure S1B and see Methods), we compared the place fields’ peak in-field firing rate differences between tracks (Track1-Track2) to the difference (Track1-Track2) in average peak instantaneous rate of those same cells during significant replay events (Figure 1C, S2). For replay events during POST, the difference in peak in-field firing rate between tracks was predictive of the difference in average peak instantaneous rates during replay events from both tracks (Figure 1D; POST: B = .03, F(1,273)= 96.8, *P* < 0.001, R2= .26; Figure S3 for individual sessions). In other words, if a place field had a higher firing rate on Track 2 (compared to Track 1), it also tended to have a higher mean firing rate across all Track 2 replay events (compared to Track 1 events) during POST. This was not the case during PRE (Figure 1D; PRE: B = .001, F(1,252)=0.12, *P* = 0.73, R2= .0004). This suggests that any contextually-driven rate modulation between two behavioral episodes is maintained during offline replay. This observation was consistent when using a more selective criteria of place field activity (e.g. place fields with higher minimum or maximum peak firing rate or place cells with a single field) (Figures S4A-B and S5D), as well as for alternative metrics for both replay firing rate (e.g. mean number of spikes per replay event and median replay rate) (Figures S5B and S5C) and place fields firing rate (e.g. place field’s area under the curve) (Figure S5A).

Replay detection methods generally detect the statistical significance of cells sequences based on templates derived from the behavioral episode, which are sensitive to the order of place field activation (during RUN and replay). However, because Bayesian decoding can also be influenced by firing rate, we next tested whether firing rate differences between tracks may bias the selection of the replay events used in the regression analysis. To remove the influence of firing rate in the replay event selection, we employed two independent types of shuffles before repeating the detection of replay events: (1) Track rate shuffle, which randomizes the overall firing rate of place fields within a track (Figures 2A–2C) and (2) Replay rate shuffle, which randomizes the cell’s firing rate during replay events (Figures 2D–2F). For both shuffles, each place cell’s firing rate was scaled based on a value randomly drawn from a distribution of firing rates obtained from the analyzed place cells on the track (for the Track rate shuffle) or during the replay events (for the Replay rate shuffle). These re-identified significant replay events were then used to repeat the regression analysis but using the original data (i.e. the original replay spike train and place fields, instead of the shuffled ones). For both shuffles, we observed a significant contextual rate modulation of replay, where track differences in the place fields’ in-peak firing rate remained predictive of track differences in the average peak instantaneous rates during replay events in POST (Figure 2C, f; POST: Track rate shuffle *P* < .001, Replay rate shuffle: *P* < .001), but not PRE (Figures 2B and 2E; PRE: Track rate shuffle *P* = .18, Replay rate shuffle: *P* = .88). As an additional control, we detected replay events using a rank order correlation method independent of Bayesian decoding (i.e. Spearman correlation), which relies only on comparing the median spike times across the place cells of a replay event compared to the sequential order of place field activity along the track (Figures 2G and 2I). Repeating the regression analysis based on the track differences in the place fields’ in-peak firing rate against the replay events detected using a Spearman correlation still resulted in a significant contextual rate modulation of the replay events rate in POST, but not PRE (Figures 2H and 2I; PRE: B = −.001, F(1,253)= .11, *P* = .74, R2= .0004, POST: B = .03, F(1,272)= 112, *P* < .001, R2= 0.3). Overall, these results indicate that the contextual rate modulation observed during replay is not a result of a bias within the replay detection analysis (namely, Bayesian decoding). Importantly, we did not observe any significant track differences in place field or replay properties (Figure S6), including the place fields’ peak in-field firing rate (Figure S6B; two-sided two-sample KS test, *P* = .59) and the replay average peak instantaneous rate (Figure S6B; two-sided two-sample KS test, *P* = .28), and confirmed that our regression analysis was not influenced by any intrinsic biases between tracks by remeasuring the statistical significance relative to a shuffled distribution (Figure S7).

**Figure 2.**
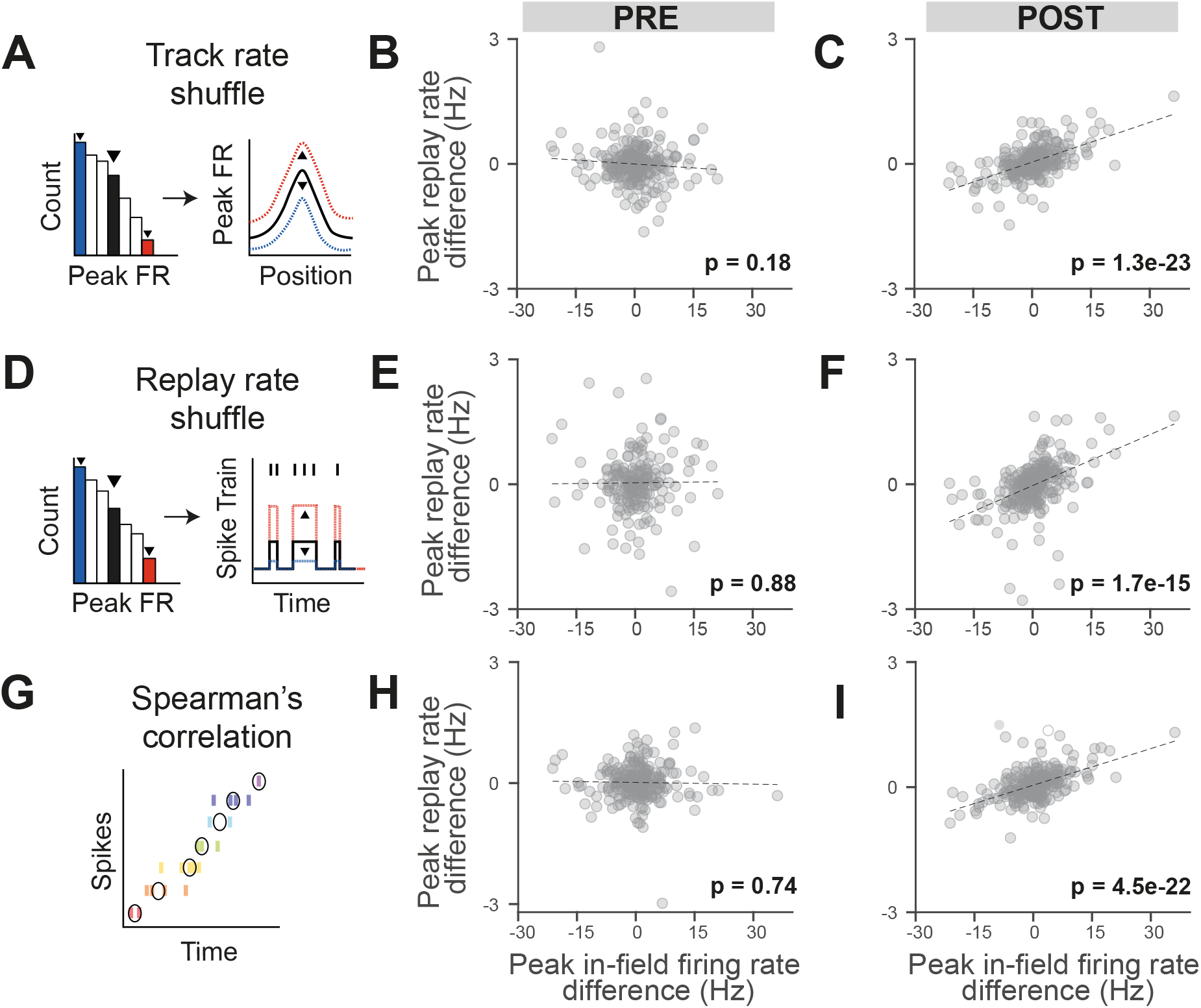
The reinstatement of rate modulation during rest replay is observed using replay detection methods insensitive to firing rate. (A, D, G) Three methods of replay detection that are insensitive to firing rate: (A) Track rate shuffle (PRE: n = 255 cells, POST: n = 274 cells), (D) Replay rate shuffle (PRE: n = 196 cells, POST: n = 263 cells), (G) Spearman correlation, where black circles indicate median spike times (PRE: n = 255 cells, POST: n = 274 cells). (A, D) Example of the original rate (right plot, in black) randomly scaled up (right plot, in red) or down (right plot, in blue), after drawing a random value from the rate distribution (left plot). (B, E, H) Regression of peak-in-field firing rate vs. peak replay rate difference (Track 1-Track 2) for PRE replay events with modified detection method: (B) Track rate shuffle, (E) Replay rate shuffle, (H) Spearman correlation. (C, F, I) Regression of peak-in-field firing rate vs. peak replay rate difference (Track 1-Track 2) for POST replay events with modified detection method (C) Track rate shuffle, (F) Replay rate shuffle, (I) Spearman correlation. The number of cells for each analysis varies as a consequence of different sets of replay events being respectively selected and therefore included in the regressions.

### Replay rate modulation emerges quickly with experience

The phenomenon of replay also occurs during awake quiescent states (e.g. while pausing on the track), and has been proposed to serve different functional roles, ranging from planning to memory storage (Diba & Buzsáki, 2007; Foster & Wilson, 2006; Gillespie et al., 2021; Pfeiffer & Foster, 2013; Xu, Baracskay, O’Neill, & Csicsvari, 2019). We observed that during the behavioral episode on the tracks, contextual rate modulation was expressed across local awake replay events (Figure 3A; RUN: B = .05, F(1,273)= 185.9, *P* < .001, R2= .41). Rate modulation during replay was even more closely aligned with behavior during RUN, compared to POST (Figure 1D; RUN: F(1,273)= 185.9, POST: F(1,273)= 96.8). Place cells during RUN already showed evidence of contextual rate modulation during awake local replay events within the first two laps of experience (Figure 3B; 2 laps: B = .06, F(1,40)= 11.85, *P* = .001, R2= .24). Furthermore, place cells expressing rate modulation during RUN replay events were also likely to show a similar direction of contextual rate modulation (e.g. Track 1 > Track 2) during POST replay events (Figure 3C; B = −.02., F(1,369)= 194.6, *P* < .001, R2= .35). While we also observed a weakly significant regression during PRE (Figure S8A; B = .03, F(1,339)= 8.73, *P* = .003, R2= .03), this regression was not significant for each of the three alternative replay detection methods that controlled for rate biases (Figure S8B-D, P>0.05). Contextual rate modulation during replay was significantly expressed throughout the entire rest period during POST (at least 1 hour cumulatively, limited only by the maximum time period tested), with the strongest effect observed in the earliest portion of POST rest (Figure 3D).

**Figure 3.**
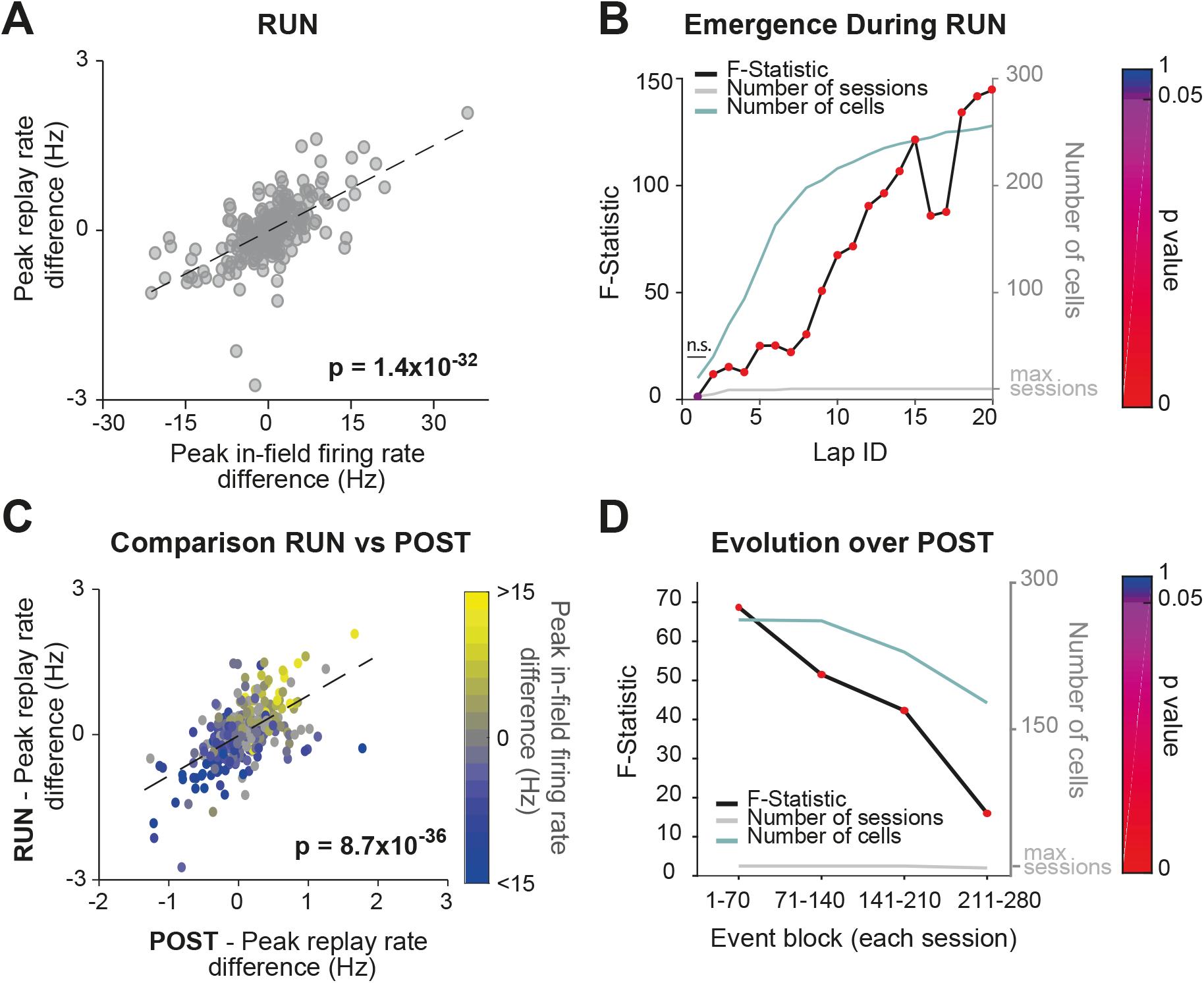
Rate modulation during replay is observed from the beginning of RUN and continues throughout the entire RUN and POST epochs. (A) Regression of peak in-field firing rates against peak replay rates (track differences) for all local RUN replay events. (B) Lap by lap emergence of rate modulation for RUN local replay events. F-statistic of regression - peak in-field firing rate vs. peak replay rate (track difference) - for replay events occurring both prior to and including the specified lap (lap ID). The p-value of each regression is indicated by color. The number of cells and number of data sessions contributing in each regression are indicated with the green and grey line, respectively. (C) Peak replay rate difference for replay events (RUN vs. POST), color coded by peak in-field rate differences between tracks. (D) Temporal dynamics of rate modulation during POST. The regression’s F-statistic (y-axis) and the corresponding p-value (marker color), analyzed in non-overlapping blocks of 70 consecutive replay events (per session). The number of cells and number of data sessions contributing in each regression are indicated with the green and grey line, respectively.

### Replay rate representations can be used to discriminate between contexts

Our data suggest that both place (i.e. sequences) and rate information is embedded within hippocampal replay events. If this is the case, then the spatial context (i.e. track identity) associated with a replay event should be identifiable using either the place or rate representations alone. To address this question, we modified the current Bayesian framework such that it would rely on either or both place and rate information (Figure 4A). For each decoded replay event, we measured the probabilistic bias towards one of the two tracks using the z-scored log odd(Carey, Tanaka, & van der Meer, 2019), 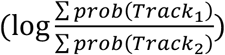. Thus, for a given replay event, this method summed the decoded posterior probability for each track, quantified the logarithmic ratio between the two tracks, and then z-scored this ratio relative to a shuffled distribution (Ratemap track label shuffle, Figure S9 and see Methods). For example, a higher positive z-scored log odd would indicate a greater probabilistic bias towards Track 1. Therefore, if the content of a replay event was detected for Track 1 (based on a statistically significant sequence), the corresponding z-scored log odd would reflect how well a place and/or rate representation could be used to correctly classify this replay event as Track 1 (rather than Track 2) (Figure 4B).

**Figure 4.**
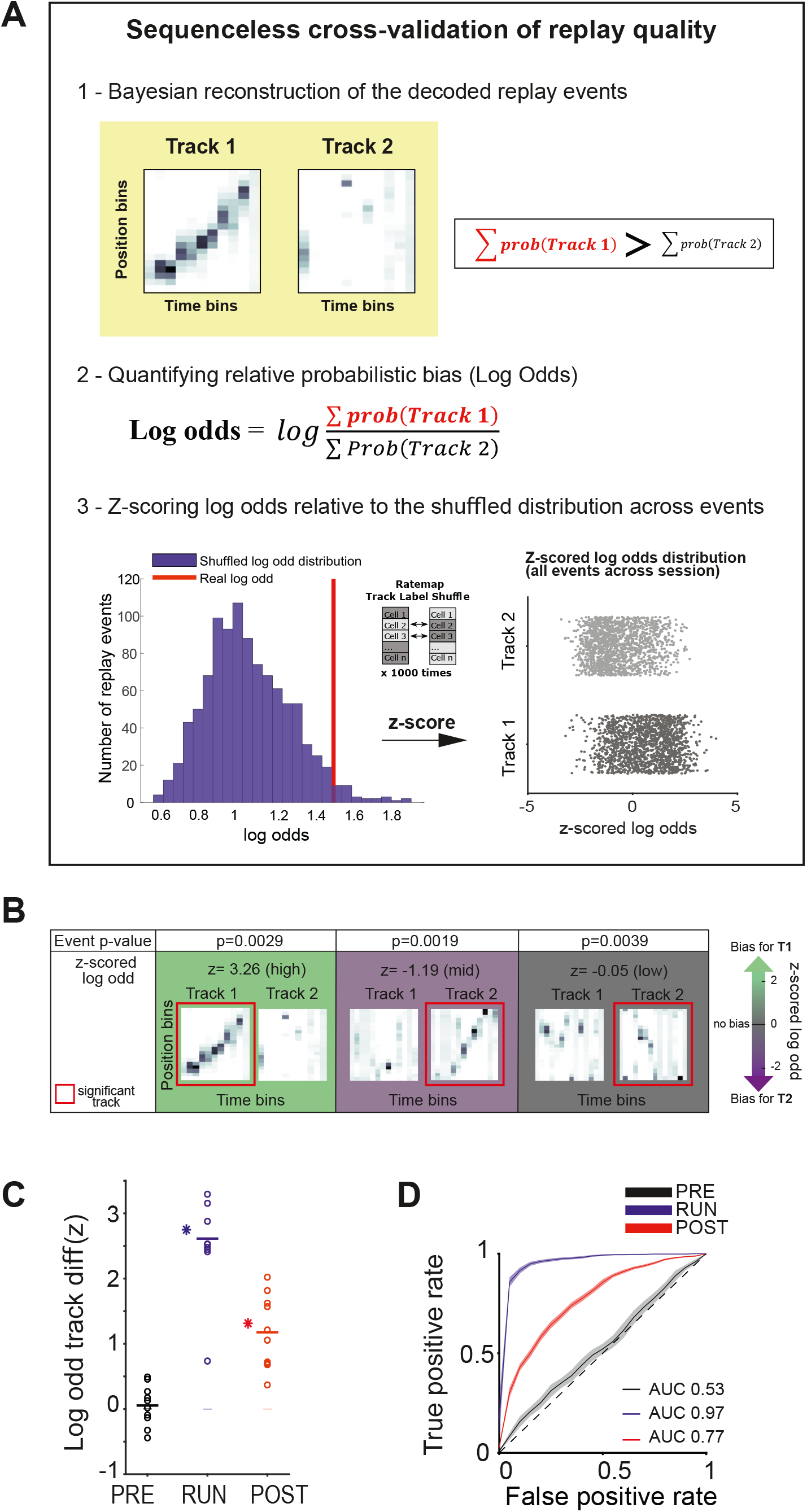
Sequenceless decoding to cross-validate replay events. (A) Description of a *sequenceless* decoding based method for cross-validating replay events originally detected based on *sequenceness*. 1) Events are decoded, and the posterior probabilities summed across each track, 2) the log odd ratio is calculated, and 3) this ratio is z-scored using a log odds distribution, based on place fields with shuffled track identity. (B) Example of three replay events scored as significant based on their sequenceness (i.e. order of cell activation) (Top row “Event p-value”), while presenting different posterior probability distributions across the two tracks (Bottom row “z-scored log odd”). From left to right, examples of replay events with high, medium and low z-scored log odd respectively. (C) The decoding performance for context (Track 1 or Track 2) using the mean z-scored log odd difference between tracks for replay during PRE (black), RUN (blue) and POST (red). Each data point is the mean of an individual session, while the horizontal line indicates the mean across all sessions (\**P* < 0.05, one-tailed rank sum test). The filled box indicates the SD from the Replay track identity shuffles. (D) ROC curves of Track1/Track2 binary discrimination for replay events during PRE, RUN and POST. The dashed line along the diagonal indicates chance performance (AUC = 0.5). The shaded region indicates the bootstrapped SE of the ROC curves.

In particular, we used two measurements to assess the replay track identity classification performance when using either the rate and/or place representation (Figure S9 and see Methods): (1) the log odd difference between two tracks compared to a Replay track identity shuffled distribution and (2) the binary discriminability of a replay event’s track identity using Receiver Operator Characteristic (ROC) curves, in which the area under the ROC curve (AUC) of 0.5 indicates chance discriminability (and an AUC of 1 indicates perfect discriminability). For both measurements, only replay events with a statistically significant replay sequence were used. When both place and rate information were unaltered, the z-scored log odd difference and the track discriminability were highest for RUN (Δ(z-scored log odd)= 2.61±.25, *P* < .001, AUC= .97), significant but marginally lower for POST (Δ(z-scored log odd)= 1.18±0.18, *P* < .001, AUC= .77), but indistinguishable from the null distribution and at chance levels for PRE (Δ(z-scored log odd)= .05±.10, *P* = .22, AUC= 0.53) (Figure 4C,D, Table S1B). Because the number of cells common to both tracks varies between sessions, and affect classification accuracy, we compared the number of cells with the mean log odd difference between tracks for each session (Figure S10). A higher number of cells was significantly predictive of a higher log odd difference for POST (B=0.34, F(1,8)= 8.34, p=.02, R2= 0.51) and RUN (B=0.055, F(1,8)= 18.97, p= .002, R2=0.70), but not PRE (B=0.011, F(1,8)= 1.84, p= .21, R2=0.19). An example of this is the presence of a session with lower log odd difference value during RUN in Figure 4C (n= 9 cells).

We then proceeded to selectively remove either the firing rate or place information available to the Bayesian decoder and assess its ability to determine the track being replayed. First, we removed the firing rate information by both setting each place field’s in-field peak firing rate to its average across both tracks and rescaling each cell’s replay spike trains to its average firing rate across all replay events (Rate fixed manipulation). We found that place information alone was sufficient to correctly classify replay events to each track, although both the z-scored log odd difference and the AUC were slightly lower than when both place and firing rate information were available (Figures 5A, E-G; AUC difference < −0.1 compared to the original for all task periods, Table S2A).

**Figure 5.**
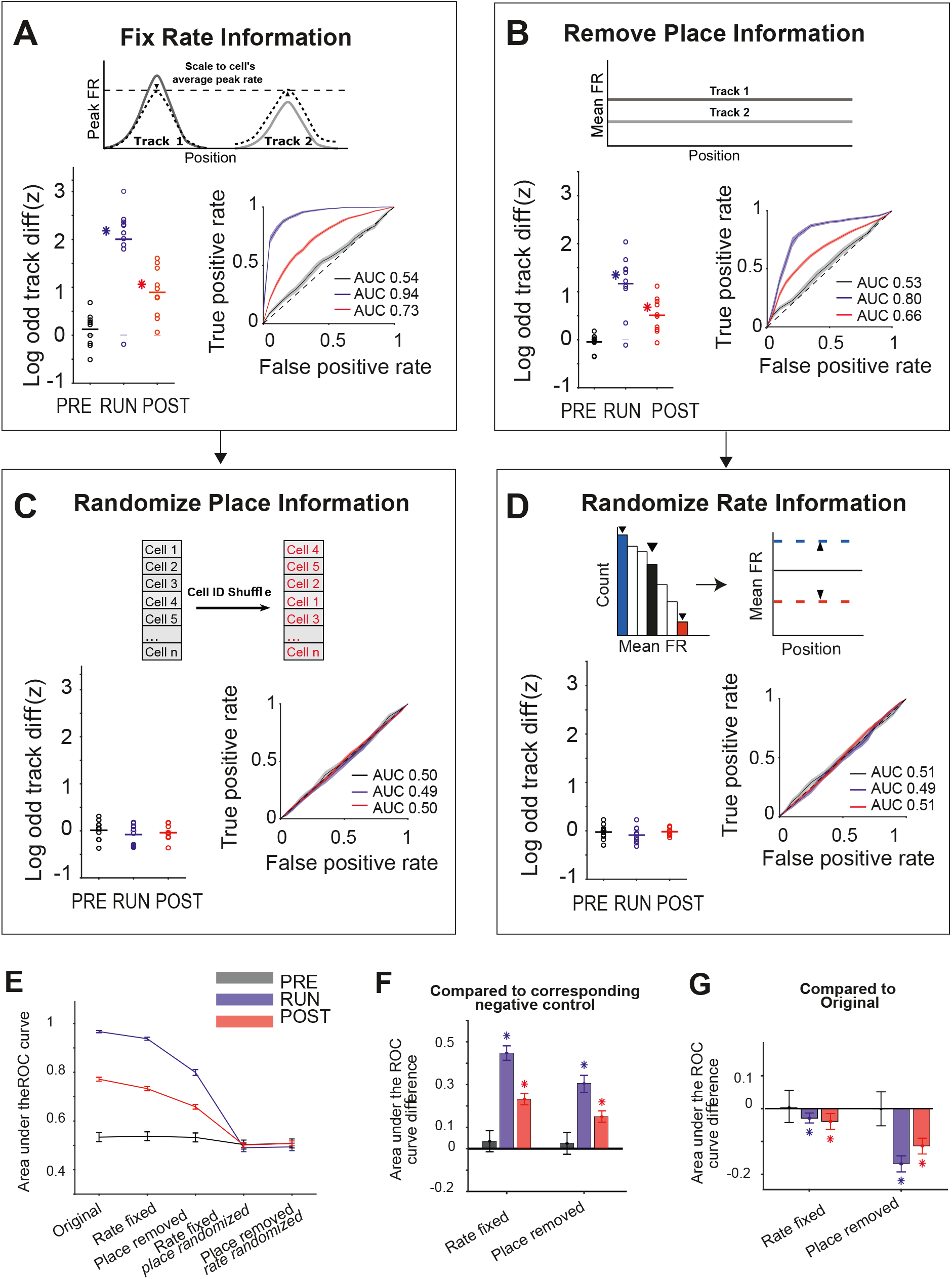
Either place or rate representations can be used to classify the replayed context. (A-D) The decoding performance for context (Track 1 or Track 2) for each manipulation, restricting the information (rate and/or place) available to the decoder. (Main plots) Mean z-scored log odd difference between tracks for replay during PRE (black), RUN (blue) and POST (red). Each data point is the mean of an individual session, while the horizontal line indicates the mean across all sessions (\**P* < 0.05, one-tailed rank sum test). The filled box indicates the SD from the Replay track identity shuffles. (Insets) ROC curves of Track1/Track2 binary discrimination for replay events during PRE, RUN and POST. The dashed line along the diagonal indicates chance performance (AUC = 0.5). The shaded region indicates the bootstrapped SE of the ROC curves. (A) Rate fixed manipulation (only place information available), (B) Place removed manipulation (only rate information available), (C) Rate fixed manipulation with place information additionally randomized (no rate or place information available), and (D) Place removed manipulation with rate information additionally randomized (no rate or place information available). (E) Summary of mean and SEM AUC under each manipulation condition, for PRE, RUN, and POST. (F) Mean AUC difference between each manipulation and the corresponding negative control. (G) Mean AUC difference between original and Rate fixed and Place removed manipulation. For (F and G), asterisks (*) indicate where the 95% confidence interval of the bootstrapped AUC difference distribution does not overlap with 0 (Tables S2A-S2B). Similar results were obtained using an alternative approach (Delong method) for measuring significance (Tables S2C-S2D and see Methods).

Next, we selectively removed place information without altering rate information, by expanding the size of both the position and time bins during the decoding: a single position bin spanning the entire track and a single time bin spanning the entire replay event (Place removed manipulation, Figures 5B and 5E-G). This manipulation still led to a significant log odd difference towards the correct track and a high track discriminability for RUN (Δ(z-scored log odds)= 1.17±.20, *P* < .001, AUC= .80), POST (Δ(z-scored log odds)= .51±.12, *P* < .001, AUC= .66), but not PRE (Δ(z-scored log odds)= −.04±0.05, *P* = 0.37, AUC= .53), supporting the finding that rate modulation is reinstated during replay and is experience-dependent. Finally, as a negative control for both manipulations, where rate and place representations were disrupted, each remaining nonmanipulated representation (place or rate) was then randomized (Rate fixed place randomized manipulation and Place removed rate randomized manipulation, respectively). Removing both place and rate representations resulted in chance level discriminability for both conditions (Figures 5C-G, Table S2B).

## DISCUSSION

Here we have shown that contextually driven place and rate representations used by the hippocampus during behavior are reinstated during replay, with either representation being sufficient to successfully discriminate which context is being replayed. These results support a model in which the hippocampus can encode more than the order of previously experienced place cell firing, with firing rates actively changing for each replay event to reflect contextual changes that have previously occurred between different behavioral episodes. This phenomenon is experience-dependent, as rate modulation within replay was only observed during and after a behavioral episode, but not prior to it. Importantly, our paradigm allows us to uniquely demonstrate that rate modulation during local awake replay emerges rapidly with experience of novel environments.

The concept that a place cell’s firing rate during a reactivation event can correlate with that of its place field in a single environment has been put forward by several previous studies (Huxter et al., 2003; Ravassard et al., 2013; Savelli & Knierim, 2019; Schwindel, Navratilova, Ali, Tatsuno, & McNaughton, 2016; Takahashi, 2015). Here, we incorporate three key advances to demonstrating the reinstatement of rate modulation. First is our use of replay, which can be identified using only the sequential pattern of firing across place cells and which contains significantly more statistical power for detection than a sequenceless based reactivation. This also avoids any circular reasoning whereby rate is used to identify the events selected for subsequent rate modulation analysis. Second, is our use of two distinct behavioral tracks, allowing us to compute the relative change in firing rates between tracks (both for behavior and replay). This avoids the problem of slow variations in neuronal excitability potentially confounding the measurement of correlation between behavior and sleep, when the analysis is limited to the absolute firing rates within a single context. It is worth noting that this study has focused on global remapping, as testing for reinstated rate modulation within a rate remapping paradigm lacks the ability to cross-reference rate changes using rate-independent replay content decoding. Finally, our approach provides a method to cross-validate replay events by comparing whether the track with the significant sequence matches the track with the higher likelihood of occupancy (log-odd analysis). This allows us to verify that both rate and place information present in replay events can independently be used downstream to identify the context being replayed.

What advantage does the reinstatement of firing rate during replay provide for memory consolidation? When minor modifications are made to an environment (e.g. changing the wall color), place cells can change their in-field firing rates without altering the location of their place fields, a phenomenon referred to as rate remapping (Leutgeb et al., 2005). As such, replay sequences would not be able to discriminate between behavioral episodes from before and after the contextual change. Our results present a possible solution to this problem, as two marginally similar contexts are still potentially separable due to their rate modulation differences. It is also important to consider that computational models of replay have commonly used modified synfire chains, sequentially connected neurons that are often embedded within distinct recurrent networks (Abeles, 1982; Chenkov, Sprekeler, & Kempter, 2017; Diesmann, Gewaltig, & Aertsen, 1999). While these models aim to replicate the sequential nature of replay, it remains to be seen whether they can also recreate experience-dependent rate modulation, or if additional mechanisms may need to be considered. Experimental evidence indicates that neocortical activity preceding a hippocampal replay event plays a key role in determining its content (Bendor & Wilson, 2012; Lewis & Bendor, 2019; Rothschild, Eban, & Frank, 2016). As such, if cortical inputs to the hippocampus during behavior underlie rate modulation (Rennó-Costa & Tort, 2017), a reinstatement of these same inputs prior to replay may be necessary for the subsequent rate modulation of hippocampal place fields during offline states.

## ACKNOWLEDGMENTS

We thank Yu Qian, Sophie Renaudineau and Julieta Campi for technical assistance; members of the Bendor Lab for valuable discussion; and Aman Saleem and Caswell Barry for their comments on the manuscript. Rat schematic in Fig. 1A adapted with permission from SciDraw.io (doi.org/10.5281/zenodo.3926277, doi.org/10.5281/zenodo.392637, https://creativecommons.org/licenses/by/4.0). This work was supported by the Biotechnology and Biological Sciences Research Council scholarship (BB/M009513/1) (MTir), the European Research Council starter grant (CHIME) (DB), the Human Frontiers Science Program Young Investigator Award (RGY0067/2016) (DB), and the Biotechnology and Biological Sciences Research Council Research grant (BB/T005475/1) (DB). The Titan Xp used for this research was donated by the NVIDIA Corporation

## AUTHOR CONTRIBUTIONS

M.Tir, M.H.G, and D.B. designed the experiment, M.Tir and M.H.G collected the data, M.Tir, M.H.G. and L.K. contributed to methodological development, M.Tir, M.H.G, and M.Tak analyzed the data, M.Tir, M.H.G, M.Tak, and D.B. wrote the initial draft, M.Tir, M.H.G, M.Tak, L.K. and D.B revised the manuscript. The order of shared first author position does not signify and should not be interpreted to suggest any difference in the work and commitment to the project by either of the individual first authors, and was determined by a coin-flip. Both MTir and MHG contributed equally and agree to reserve the right to list their name first in their CV. All authors have given approval to the final version of the manuscript.

## Methods

### Animals

Five male Lister-Hooded rats (350-450g) were implanted with a microdrive with 24 independently moveable tetrodes. Prior to surgery, rats were kept at 90% of their free-feeding weight and housed in pairs on a 12-hour light/dark cycle, with 1 hour of simulated dusk/dawn. All experimental procedures and postoperative care were approved and carried out in accordance with the UK Home Office, subject to the restrictions and provisions contained within the Animal (Scientific Procedures) Act of 1986.

### Surgery

Animals were deeply anaesthetized under isoflurane anesthesia (1.5-3% at 2L/min) and implanted with a custom-made microdrive array carrying 24 independently moveable tetrodes (modified from microdrive first published by (Davidson et al., 2009)). Each tetrode consisted of a twisted bundle of four tungsten microwires (12μm diameter, Tungsten 99.95% CS, California Fine Wire), gold-plated to reduce impedance to < 200kΩ. Three rats were implanted with a dual-hippocampal microdrive targeting both dorsal hippocampal CA1 areas (AP: −3.48mm, ML: +/- 2.4mm from Bregma), each output carrying 12 tetrodes. The two remaining rats were implanted with a microdrive targeting the right dorsal hippocampal CA1 area (AP: 3.72mm, ML: 2.5mm from Bregma) and the left primary visual cortex (AP: −5.76mm, ML: −3.8mm from Bregma), using 16 and 8 tetrodes respectively. After surgery, animals were housed individually and allowed to recover with food and water *ad libitum* for a week before returning to being kept at 90% of their free-feeding weight.

### Experimental design

A given recording session started with a 1-hour rest period in which the rats were placed in a quiet, remote location (rest pot), to which they had been previously habituated. The rest pot consisted of a black circular enclosure of 20cm of diameter, surrounded by a 50cm tall black plastic sheet that isolated them from the surroundings. The animals went through one of the two following protocols:

1. Following the rest period, the rats were exposed to two novel 2m linear tracks in which they were allowed to run back and forth for 15 min, except for one session in which the animal ran for 30 min in the second track. (Rat1 session1; Rat2 and Rat3 all sessions)
2. Following the rest period, the rats were exposed to three novel 2m linear tracks in which they were allowed to run back and forth for 15 min. Data from the first track has been removed for this study to ensure consistency between protocols, such as temporal proximity to final rest session, and for all analyses. (Rat1 session2; Rat4 and Rat5 all sessions)

Liquid reward was dispensed at each end of the track (0.1mL chocolate flavored soy milk) to encourage the animals to traverse the entirety of the track. In all except one session (Rat2 session2), the exposures to the two tracks were separated by a 10min rest period in the rest pot. The recording session finished with a final 2-hours rest period inside the rest pot.

To simulate novel environments, the shape of the tracks was changed between recording sessions and their surfaces covered with different textured fabrics. In each session, the room was surrounded by black curtains with different high contrast visual cues. The tracks were separated using view-obstructing dividers.

### Spike detection and unit isolation

Spiking data was sorted using the semi-automatic clustering software KlustaKwik 2.0 (K.Harris, http://klustakwik.sourceforge.net/) and then manually curated with either Phy-GUI (https://github.com/kwikteam/phy) or Klustaviewa (https://github.com/klusta-team/klustaviewa). Putative single units were discriminated based on the spike waveform, a clean inter-spike interval, and their stability across the recording session. The rest of the clustered activity was classified in either multi-unit activity or noise.

### LFP analysis

The power spectral density (PSD) of the LFP was calculated using Welch’s method (*pwelch*, MATLAB) to identify the channels with higher power for theta (4-12Hz) and ripple (125-300Hz) oscillations, as well as the channel with the largest difference in normalized theta to ripple power. The LFP of selected channels was next downsampled from 30 kHz to 1 kHz and band-passed filtered (MATLAB command *filtfilt*). The instantaneous phases were estimated using Hilbert transform.

### Criteria for Place cell selection

Putative principal cells were identified by selecting units with a Half Width Half Max (HWHM) larger than 500μs and mean firing rate <5Hz across the entire recording session. For place cell classification, spike trains were speed-filtered to only include the spiking activity between 4cm/s and 50cm/s. A principal cell was classified as a place cell according to the following criteria: 1) the minimum peak firing rate was >1Hz in the unsmoothed ratemap and 2) a stable spiking activity across the first half and second half of the exploration on the track. Only place cells satisfying these two criteria for each linear track independently were included in the regression analysis (Figs. 1, 2 and 3) and log odd analysis (Fig. 4,5).

To generate firing ratemaps (the spike histogram divided by the total dwell time at each position bin), the position data was discretized in 2cm bins for visualization and plotting, and 10 cm bins for Bayesian decoding. Only raw (unsmoothed) ratemaps were used for all Bayesian decoding analyses.

### Population vector analysis (PPV)

A population vector analysis was used to assess the ratemap stability for each track as well as the degree of between track remapping (Leutgeb et al., 2005). For each session, the linearized ratemaps of all recorded pyramidal cells with a firing rate > 1Hz were stacked into a 20 position bins x N cells matrix for each track. To assess remapping between tracks, the ratemaps computed from the entire track experience were compared, while to assess ratemap stability within a track, a ratemap was computed for both the first and second half of the behavioral episode. The population vector of rates at each position bin was then correlated with its counterpart vector: either on the other track or during the other half of the experience. The mean Pearson correlation value was first averaged across position bins and then across sessions.

### Bayesian Decoding

A naïve Bayesian decoding algorithm was applied to reconstruct the animal’s spatial trajectory during both behavior and replay events from CA1 hippocampal spiking activity (Zhang, Ginzburg, McNaughton, & Sejnowski, 1998):

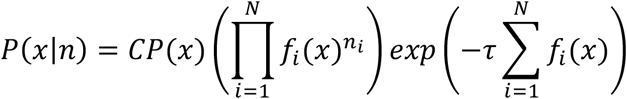

where P is the probability of the animal being at a specific position given the observed spiking activity, C is a normalization constant, x is the subject’s position, f_i_(x) is the firing rate of the i^th^ place field at a given location x, and n is the number of spikes in the time window τ. The normalization constant was re-defined as the summed posterior probabilities across both tracks. The maximum probability of the subject’s position was decoded using 250ms and 20ms time bin during behavior and replay, respectively.

The decoding error was defined as the differences between the real location of the animal and the estimated position with maximum-likelihood.

### Candidate replay event detection

Candidate replay events were detected based on MUA and ripple power. MUA was first smoothed with a Gaussian Kernel (sigma = 5 ms) and binned into 1 ms steps. Only MUA bursts with a maximum duration of 300 ms and z-scored activity over 3 were selected. Ripples LFP signal was smoothed with a 0.1 s moving average filter and a threshold over 3 was set for the z-scored ripple power.

Candidate replay events passing both thresholds were next speed-filtered (above 5 cm/s), and discarded if their sequence had less than 5 different units active or if their duration was below 100 ms or over 750 ms (thus, candidate events had at least 5 consecutive 20 ms bins). Events detected within 50 ms were combined. In an effort to optimize detection of replay events and avoid a minority of events that were discarded due to noisy probability decoding at the beginning or the end of the event, candidate replay events were split in half. To do so, the minimum MUA activity in the middle third of the candidate replay event was used to determine a natural midpoint to split the event in two segments. Both segments were decoded and tested for significance independently following the same procedure as the ‘intact’ candidate replay events (i.e., same criteria including minimum duration, etc.), for the exception of an adjusted p-value threshold (*P* < 0.025).

Replay events were classified as rest or awake local replay. Awake replay was defined as replay events occurring while the animals were running on either of the tracks and classified as local if the content of the replay event reflected the current track on which the animal was running. Replay events detected during sleep or rest periods within the rest pot were classified as rest replay events.

### Replay events scoring and significance

Replay events were scored using weighted correlation between decoded posterior probabilities across position and time.

Weighted mean:

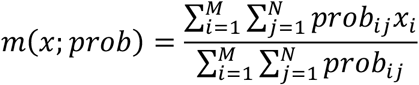

Weighted covariance:

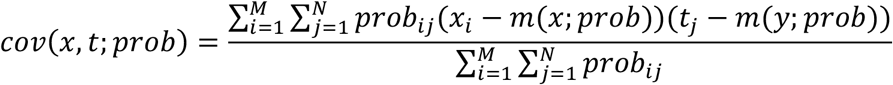

Weighted correlation:

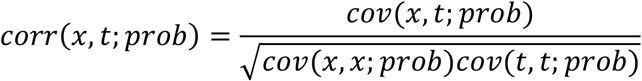

Where *x_i_*, is the i^th^ position bin, *t_j_* is the j^th^ time bin and *prob_ij_* is the probability at the position bin *i* and time bin *j*.

The statistical significance of the weighted correlation for each candidate replay event was assessed by comparison with three different 1000 shuffle distributions:

1. Spike train circular shift, in which the spike count vectors for each cell were independently circularly shifted in time within each replay event, prior to decoding.
2. Place field shift, in which each ratemap was circularly shifted in space by a random amount of position bins prior to decoding.
3. Circular shift of position, in which posterior probability vectors for each time bin were independently circularly shifted by a random amount.

If the score of the candidate event was greater than the 95th percentile of the distribution for all three shuffles then the event was considered to be significant. In a few occasions, replay events were found to be significant for both tracks. Those events were assigned to one of the tracks by computing the “Bayesian bias” score for each track. Each score was calculated as the sum of the posterior probability matrix for one track, normalized by the total sum across tracks. To assign the replay event to a specific track the Bayesian bias score had to be greater than 60%, otherwise the event was discarded.

### Reinstatement of rate modulation analysis

Linear regression was used to measure the reinstatement of rate modulation between place fields on the track and replay events (MATLAB function *fitlm*). For each place cell active on both tracks, rate modulation on the track was measured by calculating 1) the difference in peak in-field firing rate across tracks, and 2) the difference in the area under the curve (AUC; MATLAB function *trapz*). Place fields with a peak in-field firing rate < 1 Hz on either track were excluded. As a control, a variation of the analysis was done by excluding all cells with multiple place fields on the same track. Multiple peaks for a single cell were considered to be different place fields when their peak firing rate was > 1 Hz, each place field width was over 2 spatial bins (20 cm), and the distance between the peaks was of 4 spatial bins (40 cm).

Rate modulation during replay events was calculated as the change in each cell’s firing activity between the replay events encoding for each track. Included cells had to participate in at least one replay event for each track. The firing rate change was measured as 1) the difference in the average peak instantaneous rate across tracks, 2) the difference in the median rate across replay events, and 3) the difference in the mean number of spikes across replay events.

To obtain the peak instantaneous rate of each event, each cell’s spike train was binned into 1ms steps, filtered with a 100ms long Gaussian window with σ=20ms (MATLAB command *filter*), and the peak amplitude of the resulting signal measured.

### Controls for replay detection analysis

Two types of shuffles were applied to the data before repeating the detection of replay events using Bayesian decoding: a Track rate shuffle and Replay rate shuffle. The Track rate shuffle consisted in the randomization of the firing rate of the place fields within a track, while the Replay rate shuffle randomized the cells’ firing rate during the replay events. For both shuffles, each place cell’s firing rate was scaled based on a value randomly drawn from a distribution of firing rates obtained from the analyzed place cells on the track (for the Track rate shuffle) or during the replay events (for the Replay rate shuffle). The shuffled data was then used to repeat the detection of replay events, and the newly identified significant replay events were used in a linear regression comparing their original spike content (instead of the shuffled one) against the original differences in the peak in-field rate.

### Remapping classification

A bootstrapping procedure was used to identify the rate and place modulation properties of each cell. Modulation was defined when the track difference in a place field’s specific parameter (e.g. peak in-field firing rate or peak location) exceeded the intrinsic variability of such parameter on a single track. The distribution of each parameter was computed for both tracks by calculating firing ratemaps from a randomly sampled N out of N laps (1000 iterations, with replacement). The median value of a track distribution was then compared to the 5^th^/95^th^ percentiles of the other track, and vice versa. If either median exceeded either percentile, the cell was classified as being modulated between tracks for that parameter.

### Sequenceless Bayesian Decoding

The sequenceless Bayesian decoding approach used here was modified from a previous method described by Carey et al., 2019. Only cells with stable place fields on the track in the first and second half of the behavioral episode (peak in-field firing rate > 1Hz) and replay events significant for a single track were included in our sequenceless decoding analysis.

Before Bayesian decoding, ratemaps for Track 1 and Track 2 were concatenated as a single matrix [nCell x (nPosition(T1) + nPositions(T2))]. Unless otherwise mentioned, 20ms time bins and 10cm position bins were used for decoding. After decoding, the posterior probabilities for each time bin were normalized across all position bins from both tracks to sum to 1. Then, the summed posterior probability across all time bins within each replay event was computed, and the probabilistic bias towards Track 1 or Track 2 for each replay event was quantified as 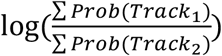 (log odds), where ∑*Prob*(*Track*_1_) and ∑*Prob*(*Track*_2_) referred to the summed posterior probabilities for Track 1 and Track 2, respectively. To remove any intrinsic probabilistic bias, each raw log odd calculation was then z-scored relative to a shuffled log odd distribution (Ratemap track label shuffle), obtained by randomly permuting each cell’s Track 1 and Track 2 ratemaps 1000 times for each replay event. The permutation was done by either keeping a cell’s ratemap unaltered or performing a between track swap.

To test the decoding when only using place or rate information, the original data were manipulated as the following:

- Rate fixed manipulation: Rate information was fixed both by rescaling each cell’s peak infield firing rate to the mean value across both tracks, and by rescaling its spike count in each replay event to the average firing rate across all replay events.
- Place removed manipulation: The place (and sequence) information was removed by decoding with only a single position bin, which encompassed the entire track (mean firing rate on each track when speed is above 5cm/s) and a single time bin, which spanned the entire replay event.

Two negative controls were used to disrupt both rate and place information:

- Rate fixed place randomized manipulation: After the Rate fixed manipulation, place information was randomized using a cell ID shuffle in which all cells were randomly reassigned to a different cell’s ratemap for each replay event.
- Place removed rate randomized manipulation: After Place removed manipulation, rate information was randomized by scaling the firing rate on each track based on a value randomly drawn from a distribution of firing rates obtained from the analyzed place cells on the track.

Given the inherent random nature of our shuffling procedure, it was important to ensure that the result of a single simulation was reproducible. Thus, for both negative control manipulations, we ran our simulations 10 times. We observed that 9/10 times the results were not significant (ranksum test, P>0.05, uncorrected). The data presented in Fig. 5 was selected from the negative control simulation nearest the median log-odd difference (out of 10 simulations) (Table S3).

### ROC analysis

The ‘ground truth’ track identity for each replay event was assigned based on the sequenceness of the replay, which was determined by comparing the event’s weighted correlation score to three shuffled distributions. The binary discriminability of a replay event’s identity (Track 1 or Track 2) during PRE, POST or RUN was quantified using the Receiver Operative Characteristic (ROC) curve. To ROC curve was constructed based on a range of true and false positive rates obtained by systematically shifting the discrimination threshold along the z-scored log odd distributions, for both Track 1 and Track 2 replay events. A trapezoidal approximation was used to estimate the area under each ROC curve (MATLAB function *perfcurve* for both constructing ROC curve and estimating area under the curve). The bootstrapped distributions of ROC curves and the area under the corresponding ROC curves were produced by resampling with replacement 1000 times.

### Statistics

#### Z-scored Log Odd significance

The statistical significance of the mean difference between Track 1 and Track 2 z-scored log odd is quantified using a one-tailed rank sum test in which mean z-scored log odd track differences across sessions was compared to the mean shuffled differences across sessions. The shuffled difference distribution for each session was obtained by resampling replay events 1000 times with replacement and assigning their track identities randomly (referred to as Replay Track Identity Shuffle). The mean z-scored log odd difference was considered significantly different from the shuffled distribution at the significance level of 0.05.

#### Bootstrapping for ROC significance

To determine if rate and/or place representation manipulations significantly changed the area under the curve (AUC) of the ROC curves, we calculated the confidence interval for the difference between each condition’s bootstrapped AUC distribution and that of the original data. This was repeated in an additional analysis by replacing the original data by one of the negative controls (Rate fixed place randomized manipulation and Place removed rate randomized manipulation). The mean difference between two bootstrapped AUC distributions were only considered statistically significant when the 95% confidence interval did not overlap with 0.

#### Delong Test for ROC

To determine if rate and/or place manipulations statistically significantly changed the AUC of ROC curves within the same behavioral epochs, the AUC of both manipulations (i.e. Rate fixed or Place removed) were compared with the AUC of the original data or their corresponding negative controls (i.e. Rate fixed Vs Rate fixed place randomized and Place removed Vs Place removed rate randomized). The DeLong Test (Sun & Xu, 2014) was employed to perform pairwise comparisons between two ROC curves. The DeLong variance-covariance matrix for two ROC curves was computed using MATLAB-based algorithm [https://github.com/PamixSun/DeLongUI] developed by Sun and Xu (2014)(Sun & Xu, 2014). A z-score was calculated based on the following equation:

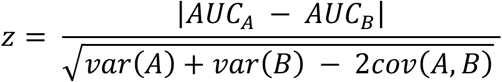

where AUC referred to the area under the ROC curve, var(X) referred to the variance of ROC curves from a 1000 bootstrapped distribution, and cov(X) referred to the covariance of ROC curves from both 1000 bootstrapped distributions.

The z-score for each pair of comparisons was subsequently used to calculate the two-tailed p-value. Two AUC distributions were considered significantly different when *P* ≤ 0.05.

All analysis was carried out using MATLAB (Mathworks, Natick MA).

## Data availability

All data used in this manuscript are available from the authors upon reasonable request.

## Code availability

All custom-written code is available upon request.

