## Supplemental Figures for "Experience-Driven Rate Modulation is Reinstated During Hippocampal Replay"

### SUPPLEMENTAL INFORMATION

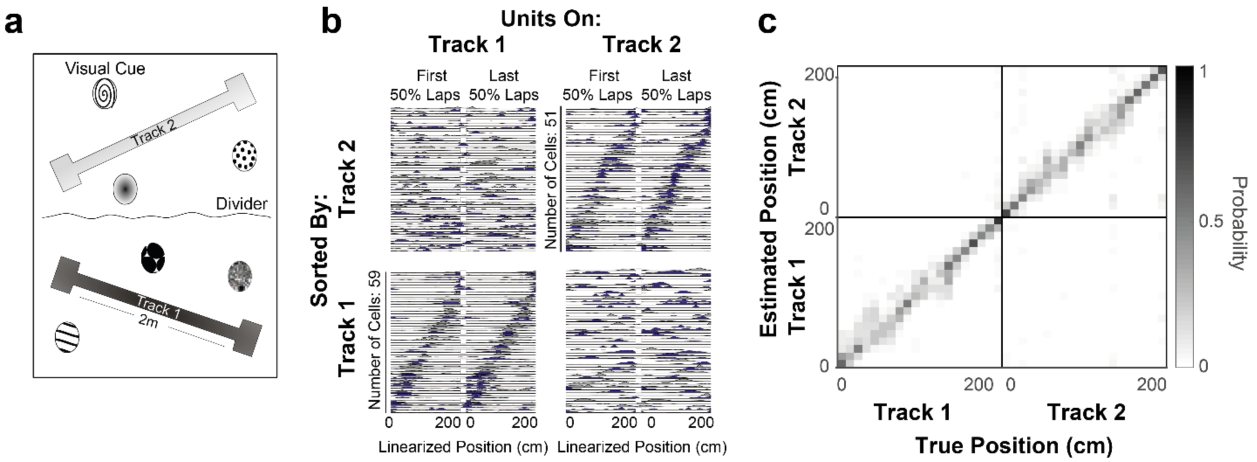

**Figure S1. Experimental setup, remapping and field stability.**

(A) Layout of the recording room - two linear tracks, each surrounded by visual cues and visually occluded from each other.

(B) Ratemaps for Track 1 (left column) and Track 2 (right column) in an example session (rat 3, session 2) computed for the first 50% of laps (left half) and last 50% of laps (right half) and sorted by the ordered position of place fields on Track 1 (bottom) and Track 2 (top).

(C) Confusion matrix for decoded position (posterior probability normalized over both tracks) for an example session (rat 3, session 2). Distribution of peak posterior probabilities (y-axis) is plotted at each True position during the behavioral episode (x-axis).

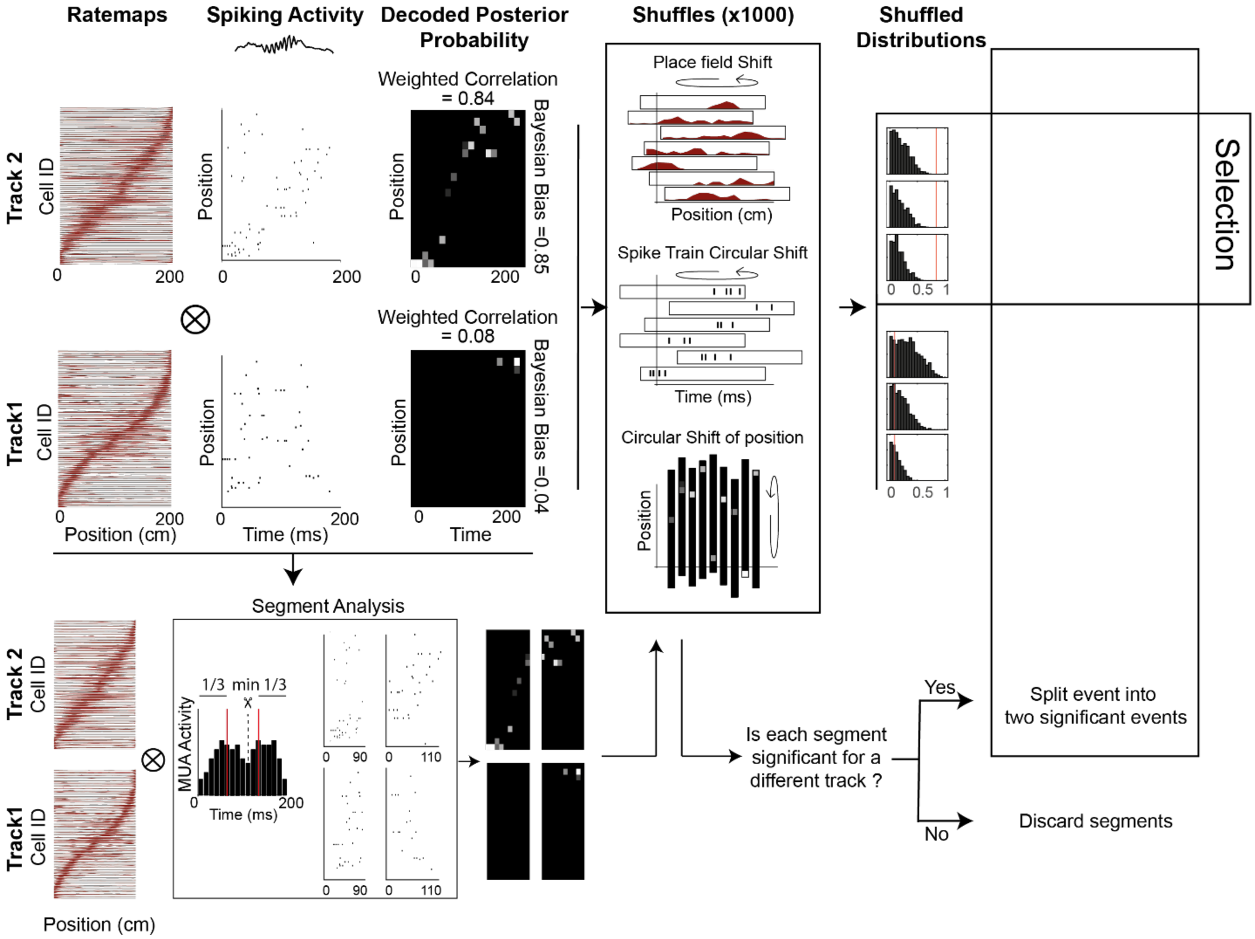

**Figure S2. Analysis pipeline for detection of significant replay events.**
Detected replay spike trains were decoded using a Bayesian framework. Each track's ratemaps obtained during RUN were used to calculate the decoded posterior probabilities for each replay sequence (top left). In addition, all replay events were split into two shorter events based on the minimum in the MUA during the middle third of the replay event (bottom left, Segment analysis). For each replay event, weighted correlations were calculated on decoded posterior probability from both the whole replay sequence and each segment, and the three scores were compared to three different shuffles (Place field shift, Spike train circular shift, and Circular shift of position, 1000 shuffles performed for each type). Only events with weighted correlation scores higher than 95% for every shuffle distribution (97.5% for split replay events) were considered statistically significant.

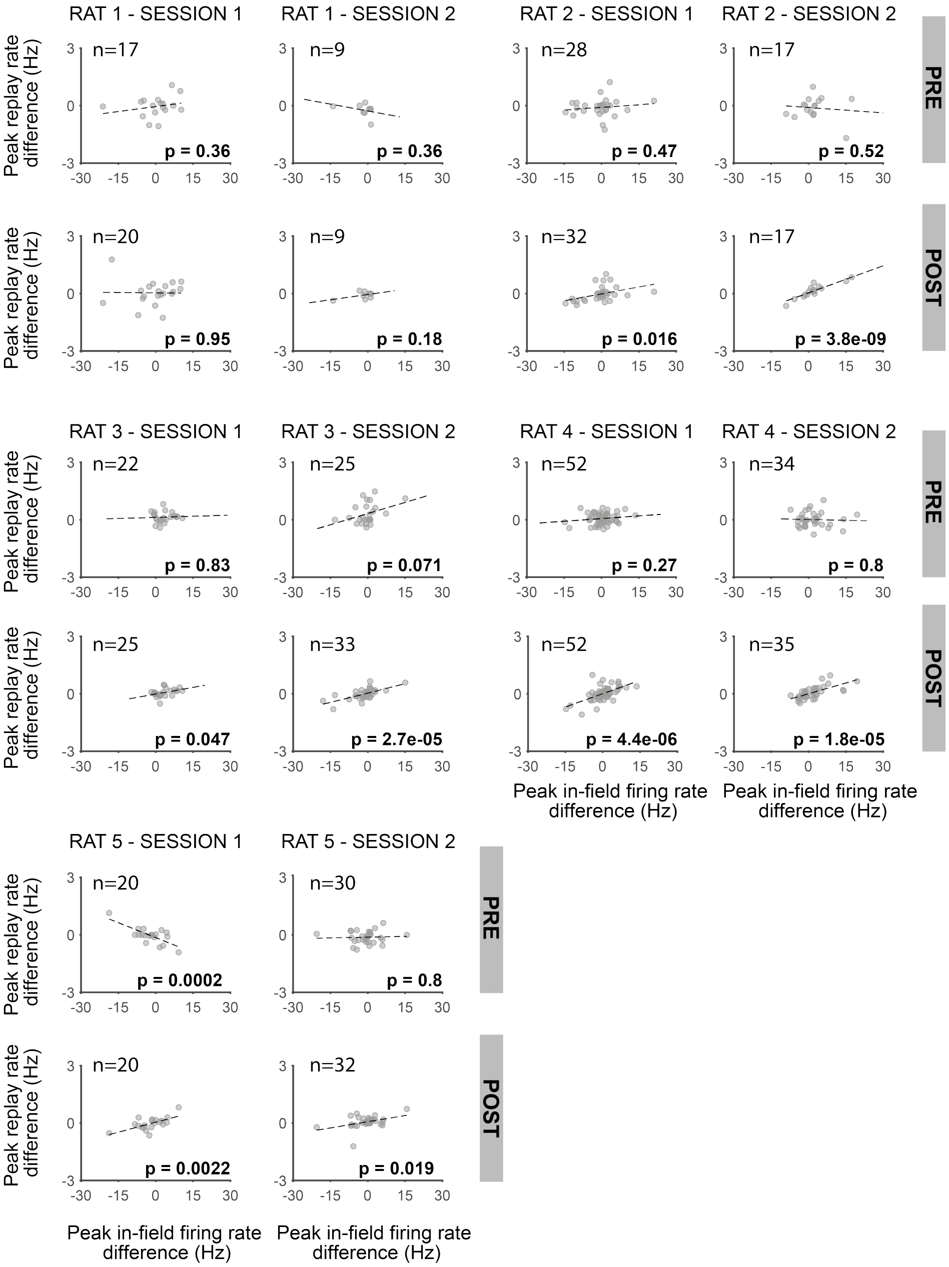

**Figure S3. Reinstatement of rate modulation during replay for each individual session.**

The peak in-field firing rate difference (Track1-Track2) significantly predicts the average peak instantaneous replay rate difference (Track1-Track2) in POST in eight out of ten session. In contrast, only one out of ten session have a statistically significant regression in PRE. Each data point is a neuron active on both tracks. Significance and number of data points (common cells in both tracks) are indicated in each plot.

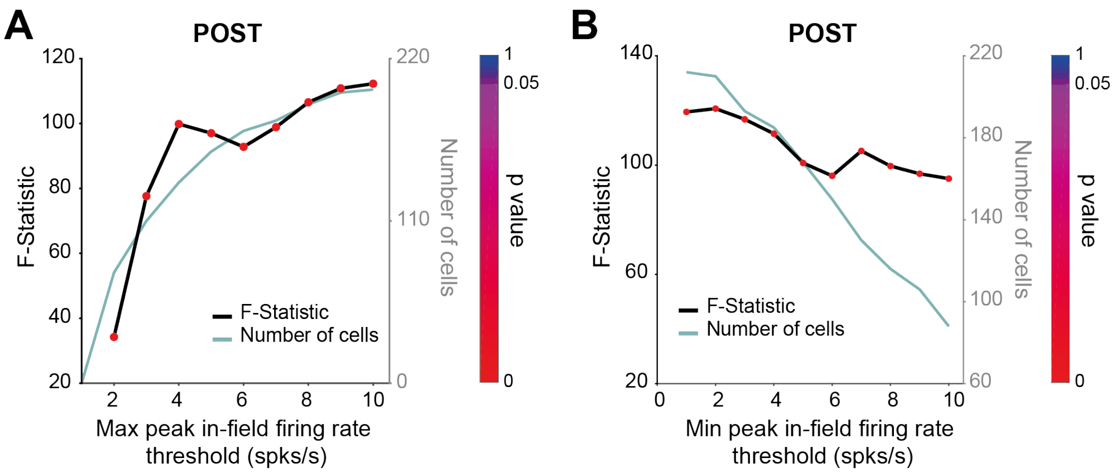

**Figure S4. Significance of regression analysis using a stricter criterion of place cell activity.**

**(A)** F-statistic of regression - peak in-field firing rate vs. peak replay rate (track difference) - for replay events occurring during POST, as the criterion for place cell activity (maximum peak in-field firing rate) is progressively increased from 1 Hz to 10 Hz. The first regression which only included place cells with a maximum peak-in-field firing rate of 1Hz is absent as no cells met this criterion. All the remaining regressions are statistically significant (up to and including using a threshold of 10 spk/s). The p-value of each regression is indicated by color. The number of cells used in each regression is indicated by the green line.

**(B)** F-statistic of regression - peak in-field firing rate vs. peak replay rate (track difference) - for replay events occurring during POST, as the criterion for place cell activity (minimum peak in-field firing rate) is progressively increased from 1 Hz to 10 Hz. All regressions are statistically significant (up to and including using a threshold of 10 spk/s). The p-value of each regression is indicated by color. The number of cells used in each regression is indicated by the green line.

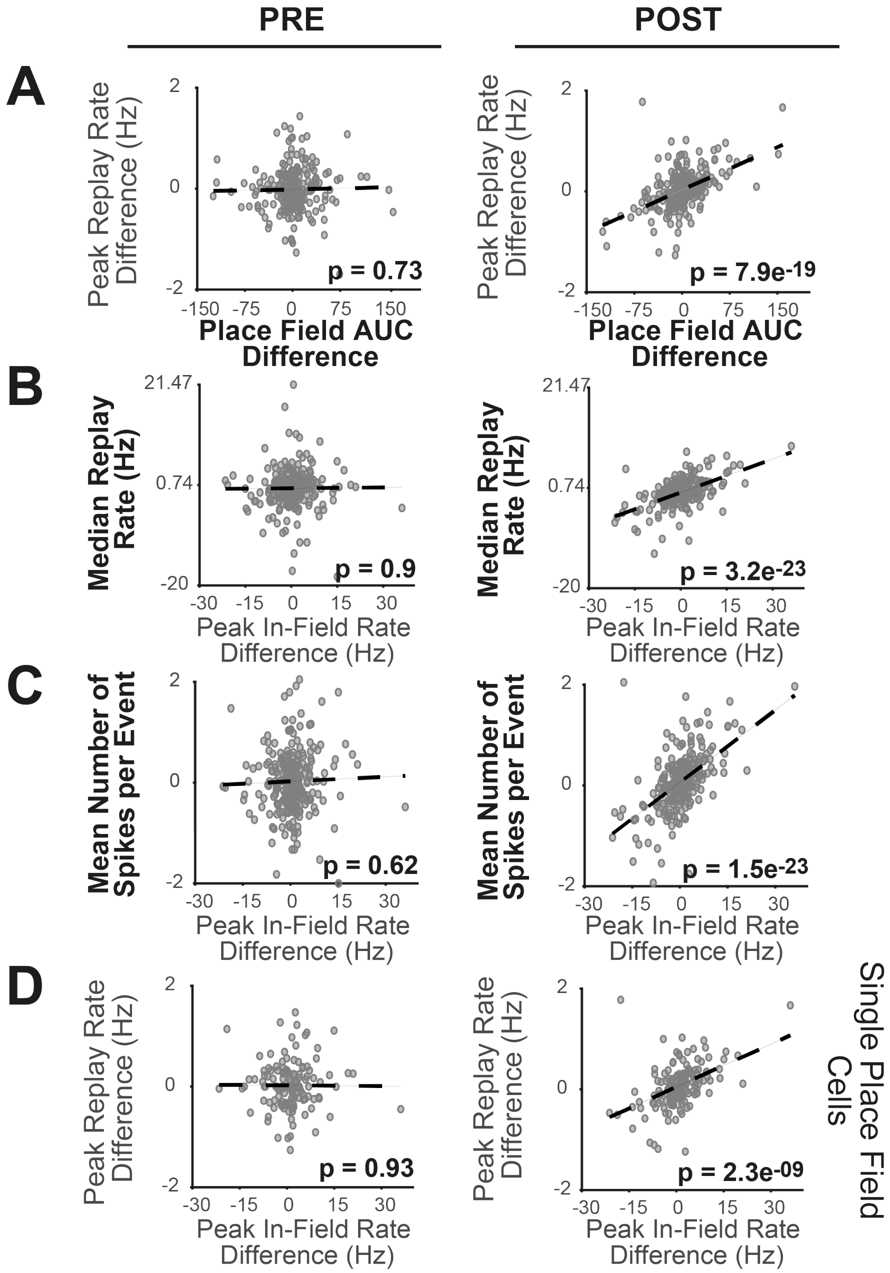

**Figure S5. Significance of regression analysis with alternative place field and replay rate metrics, and stricter criteria for cell selection.** F-statistic of regression - peak in-field firing rate vs. peak replay rate (track difference) - for replay events occurring during PRE (left column) and POST (right column) using alternative metrics: (A) Place field area under the curve (new x-axis, PRE: n = 254 cells, POST: n = 275 cells), (B) Median replay rate (new y-axis, PRE: n = 254 cells, POST: n = 275 cells), and (C) mean number of spike per event (new y-axis, PRE: n = 254 cells, POST: n = 275 cells). And using new criteria for cell selection: (D) Single place field cells (PRE: n = 138 cells, POST: n = 155 cells).

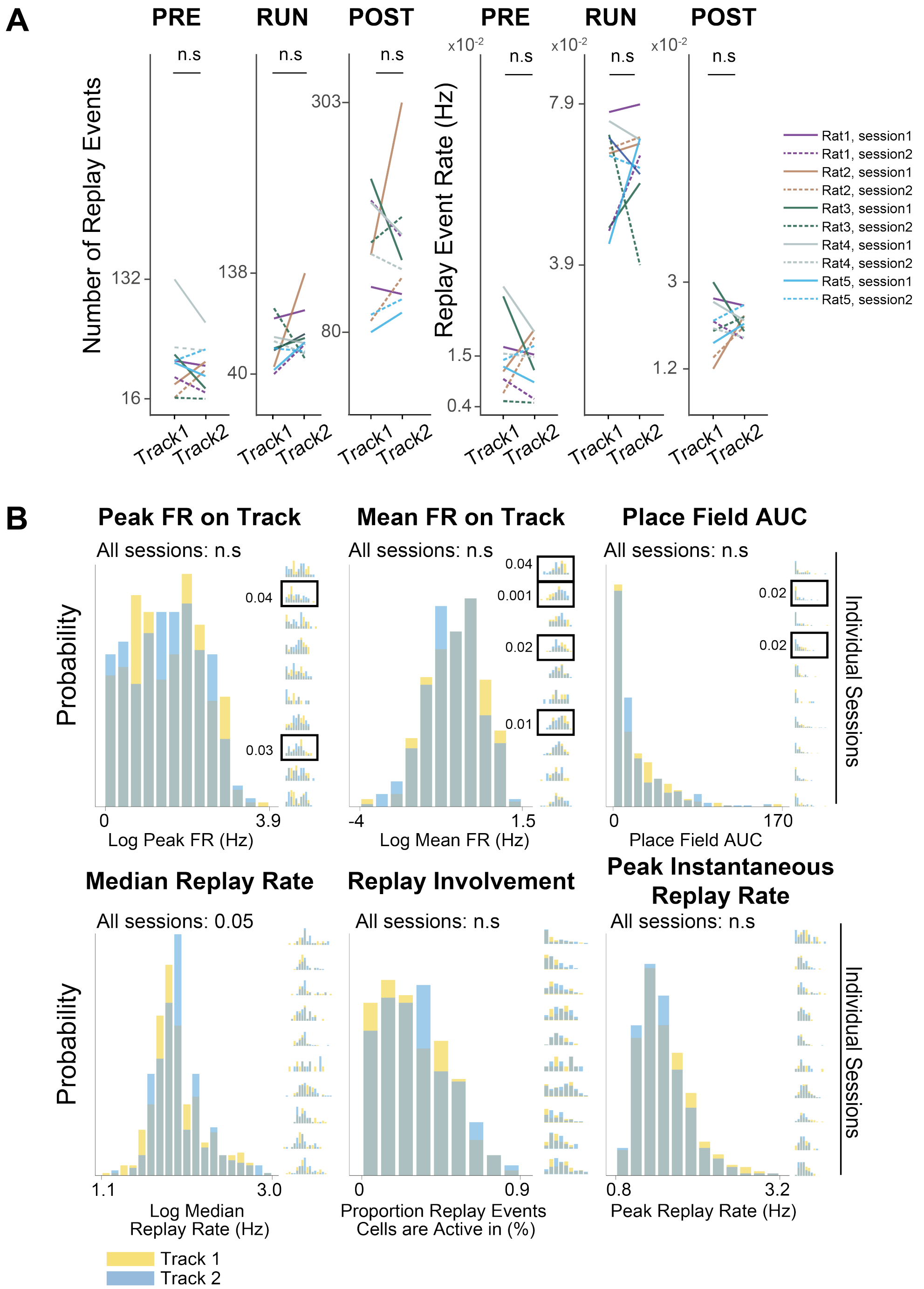

**Figure S6. Between track comparison of place field and replay properties.**

(A) Number of replay events (left) and replay event rate (right) for each track during PRE, RUN, and POST. The differences between tracks for any of these comparisons were not statistically significant (Wilcoxon sign rank, $P > 0.05$, n=10).

(B) Distribution of Peak firing rate on track (top left), mean firing rate on track (top center), place field AUC (top right), median replay rate (bottom left), replay involvement (bottom center) and peak instantaneous replay rate (bottom right) for tracks 1 and 2, and for individual session distributions (inset) (K-S test, n.s. $P > 0.05$).

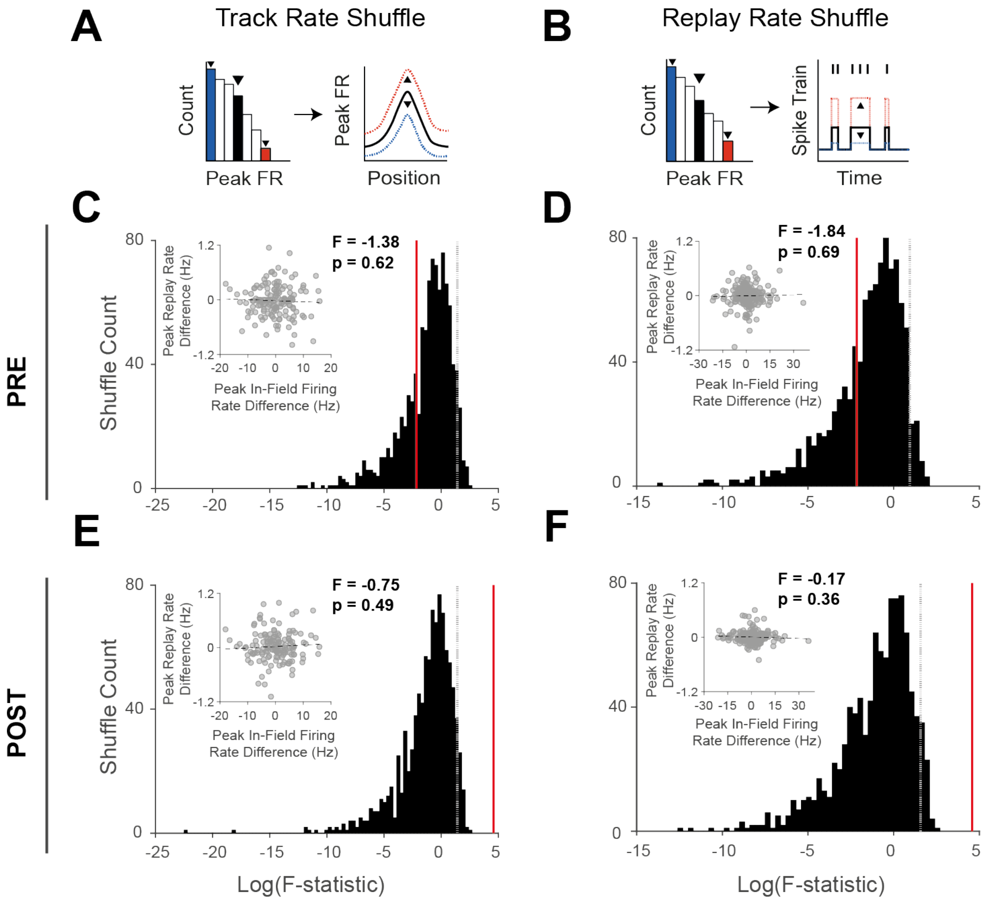

**Figure S7. Re-computing the statistical significance of regression analysis relative to a shuffled distribution.**

(A) Track rate shuffle and (B) Replay rate shuffle were used to randomize the overall firing rate of place fields within a track (x-axis) and the cell's firing rate during replay events (y-axis), respectively. A F-Statistic distribution was then calculated to measure the significance of the regression. (A and B) Example of the original rate (right plot, in black) randomly scaled up (right plot, in red) or down (right plot, in blue), after drawing a random value from the rate distribution (left plot).

(C to F) F-statistic distributions obtained from the shuffle (black), where dashed gray line indicates $P < 0.05$, and red line indicates the F-Statistic value of original data (main plot Figure 1D). One example of a regression plot within the shuffle distribution (inset plot). (C) PRE: Track rate shuffle. (D) PRE: Replay rate shuffle. (E) POST: Track rate shuffle. (F) POST: Replay rate shuffle.

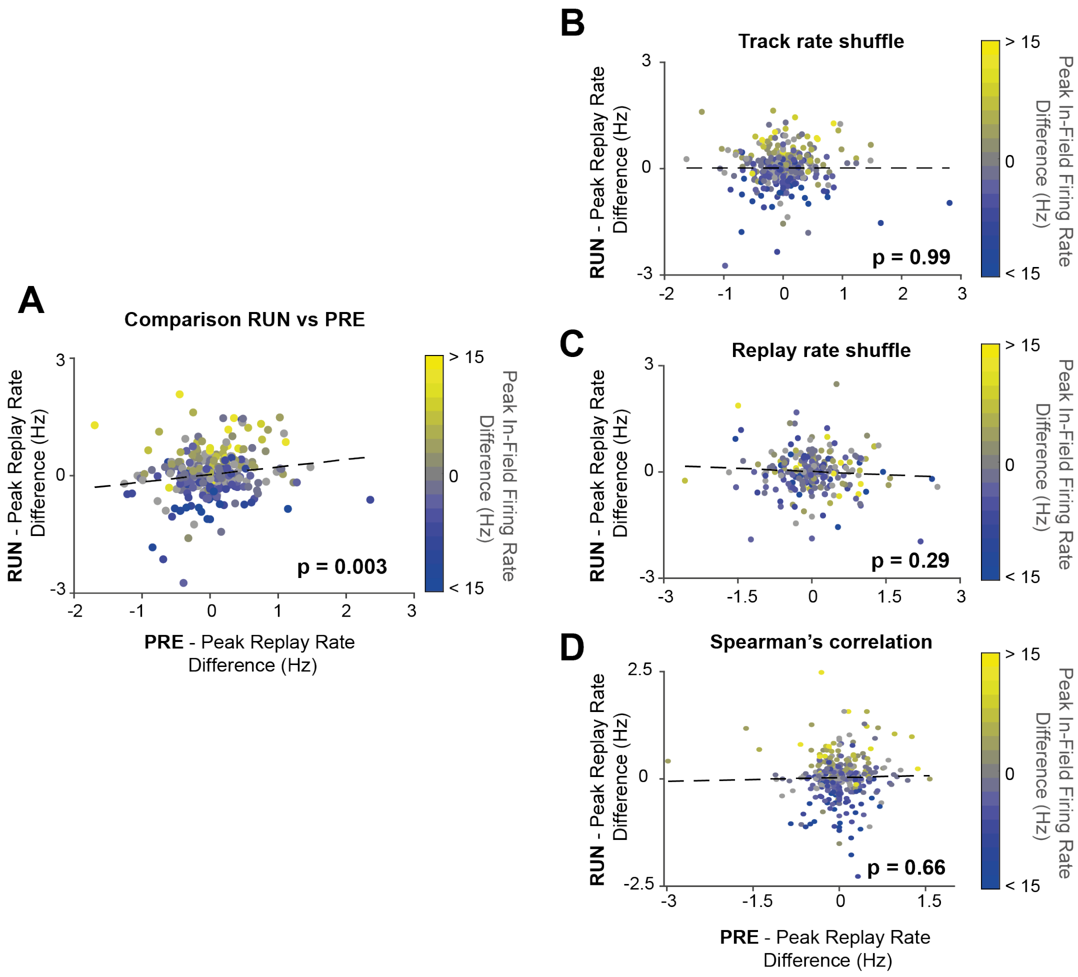

**Figure S8. Between track differences in replay rate during PRE is not predictive of replay rate differences observed during RUN local replay events.** Peak replay rate difference for replay events (RUN vs. PRE), color coded by peak in-field rate differences. A linear regression was weakly statistically significant (*P* =0.003) using our main replay event detection method, however we did not observe a significant regression. (*P* > 0.05) for any of the three alternative methods that control for rate biases in selective significant replay events– B. Track rate shuffle, C. Replay Rate shuffle, and D. Spearman correlation.

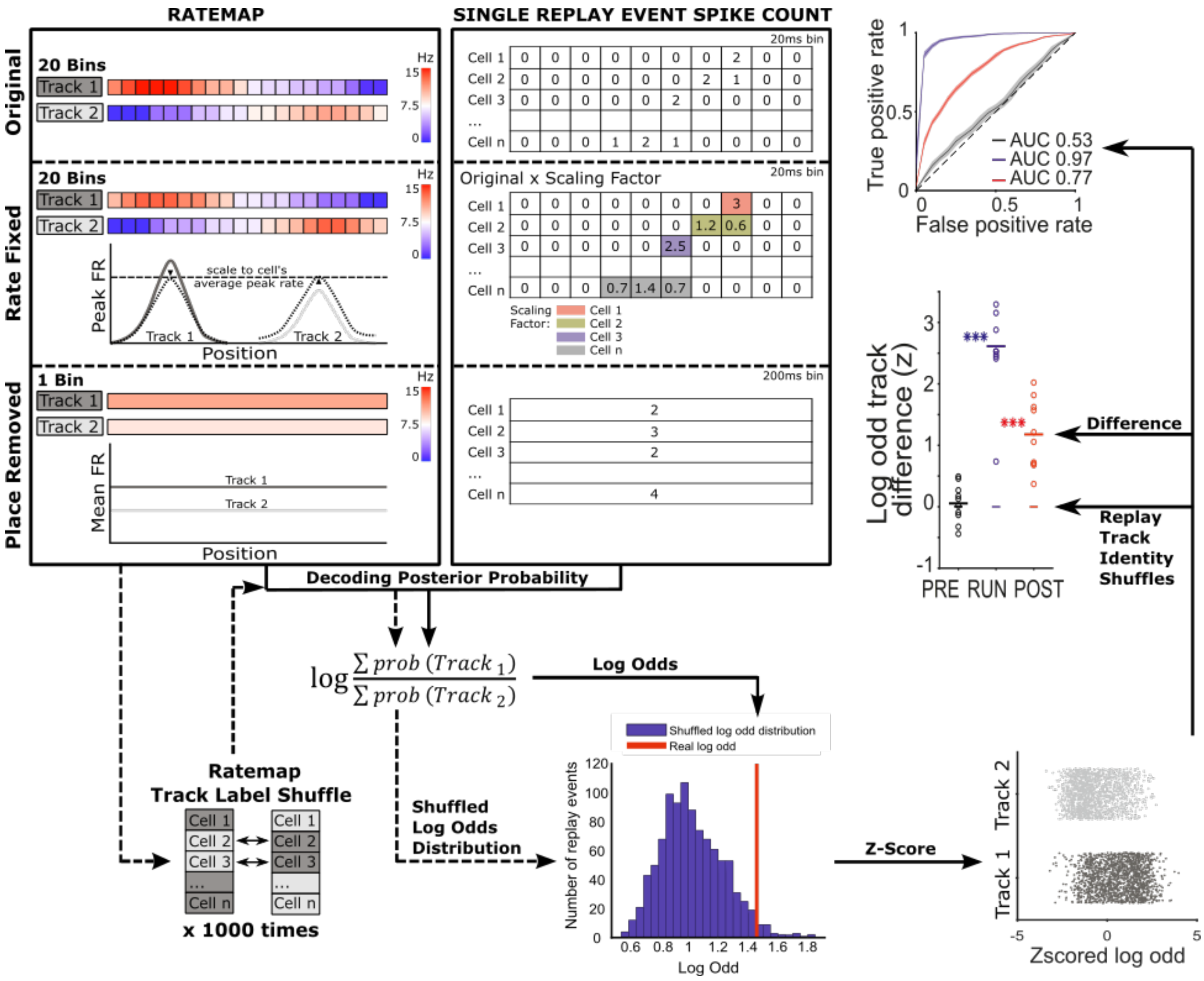

**Figure S9. Schematic of modified Bayesian decoding framework.** The concatenated Track 1 and Track 2 ratemap (top first column) and spike counts during each replay event (top second column) are used for Bayesian decoding. This produces a posterior probabilistic distribution for each replay event, which is normalised across all position bins from both tracks to sum to 1 (Original, top row). Then, the relative probabilistic bias between Track 1 and Track 2 (Log odd) is computed. As control conditions, the original ratemap and replay spike counts are selectively modified such that only place information (Rate fixed manipulation, second row) or rate information (Place removed manipulation, third row) is available for Bayesian decoding. For Rate fixed manipulation, rate information is fixed by 1) rescaling each cell's peak in-field firing rate to the mean value across both tracks, and 2) rescaling its spike count during each replay event to the average firing rate across all replay events. For Place removed manipulation, the place and sequence information is removed by decoding using only a single position bin covering the entire track (when the animal's moving speed is above 5cm/s) and a single time bin spanning the entire replay event. For all conditions, the manipulated replay events are next decoded using the original decoding pipeline, including a Track 1 and Track 2 Ratemap Track label shuffle, obtained by randomly permuting each cell's Track 1 and Track 2 ratemaps 1000 times for each replay event (bottom left). The permutation is done by either keeping a cell's ratemap unaltered or performing a between track swap. The z-scored log odd for each replay event is obtained by comparing the raw log odd relative to the shuffled log odd distribution. For each behavioral epoch (i.e PRE, RUN and POST), Track 1 and Track 2 z-scored log odd distributions are obtained by classifying each replay event according to the track identity determined by the sequenceness of the replay (bottom right). The track discrimination performance is quantified using two measurements.

The first method quantifies binary discriminability using receiver operative characteristic curve (ROC curve). In brief, the ROC curve is constructed by plotting the true positive rate against false positive rates obtained by shifting discrimination threshold across the two z-scored log odd distributions. The area under the ROC curve (AUC) would thereby indicate the performance of binary discriminability where AUC = 1 indicates perfect discrimination and AUC = 0.5 indicates chance level discrimination (right, top plot). The second method quantifies the mean difference between Track 1 and Track 2 z-scored log odd distributions. The mean z-scored log odd track difference for each behavioural epoch is obtained by taking the average of the z-scored log odd track difference across all replay events within each session and then across all ten sessions. The statistical significance for this measurement is quantified using one-tailed rank sum test in which the mean z-scored log odd track difference from 10 sessions are compared to the shuffled version of the log odd track differences distribution where the replay events from each session were resampled with replacement 1000 times and their track identities were assigned randomly (referred to as Replay Track Identity Shuffle) (right, bottom plot).

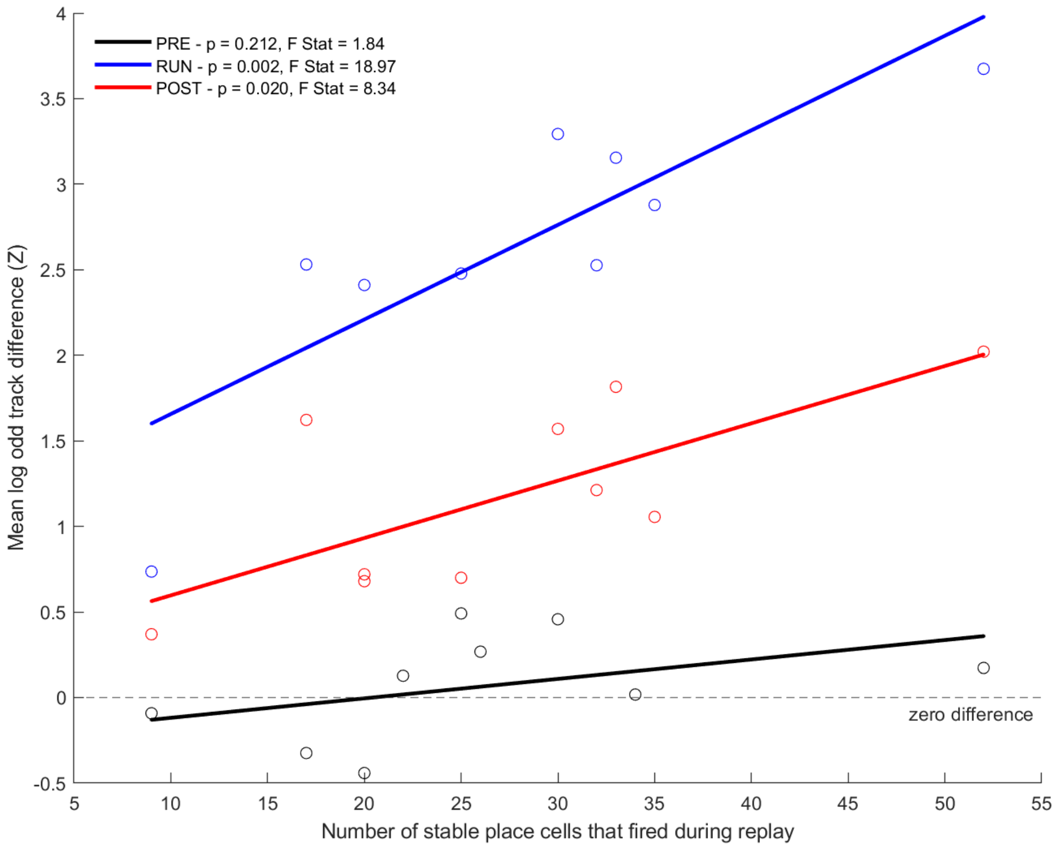

Figure S10. **The relationship between the number of stable place cells that participated in replay and the mean log odd track difference during PRE (Black), RUN (Blue) and POST (Red).** Each dot represented the mean value of an individual session. The horizontal dashed line indicated zero track difference. A linear regression was considered statistically significant when $P < 0.05$.

| **a** | | | Time resting (min) | | Cells | | | Decoding quality (%) | |
|---|---|---|---|---|---|---|---|---|---|
| Session | Track | Number of laps | PRE | POST | Cells used for replay analysis | common to T1 and T2 (stable/non-stable)* | Included in regression analysis | Classification accuracy | Local decoding accuracy |
| Rat 1, session1 | 1 | 30 | 72 | 76 | 42 | 30 (20/10) | 20 | 91 | 81 |
| Rat 1, session1 | 2 | 20 | 72 | 76 | 44 | 30 (20/10) | 20 | 97 | 82 |
| Rat1, session2 | 1 | 10 | 83 | 155 | 27 | 10 (9/1) | 9 | 86 | 58 |
| Rat1, session2 | 2 | 12 | 83 | 155 | 27 | 10 (9/1) | 9 | 93 | 66 |
| Rat 2, session 1 | 1 | 15 | 59 | 125 | 70 | 37 (30/7) | 30 | 87 | 86 |
| Rat 2, session 1 | 2 | 59 | 59 | 125 | 63 | 37 (30/7) | 30 | 80 | 89 |
| Rat 2, session 2 | 1 | 22 | 54 | 104 | 49 | 28 (17/11) | 17 | 92 | 82 |
| Rat 2, session 2 | 2 | 33 | 54 | 104 | 43 | 28 (17/11) | 17 | 90 | 79 |
| Rat 3, session 1 | 1 | 34 | 49 | 123 | 59 | 33 (25/8) | 25 | 94 | 85 |
| Rat 3, session 1 | 2 | 27 | 49 | 123 | 51 | 33 (25/8) | 25 | 94 | 80 |
| Rat 3, session 2 | 1 | 36 | 69 | 137 | 72 | 44 (33/11) | 33 | 94 | 85 |
| Rat 3, session 2 | 2 | 38 | 69 | 137 | 57 | 44 (33/11) | 33 | 97 | 86 |
| Rat4, session1 | 1 | 19 | 103 | 129 | 87 | 66 (52/14) | 52 | 94 | 80 |
| Rat4, session1 | 2 | 10 | 103 | 129 | 80 | 66 (52/14) | 52 | 94 | 85 |
| Rat4, session2 | 1 | 13 | 98 | 125 | 63 | 44 (35/9) | 35 | 88 | 78 |
| Rat4, session2 | 2 | 11 | 98 | 125 | 52 | 44 (35/9) | 35 | 97 | 82 |
| Rat5, session1 | 1 | 13 | 91 | 76 | 50 | 31 (20/11) | 20 | 97 | 85 |
| Rat5, session1 | 2 | 11 | 91 | 76 | 57 | 31 (20/11) | 20 | 90 | 89 |
| Rat5, session2 | 1 | 13 | 72 | 86 | 70 | 47 (32/15) | 32 | 94 | 88 |
| Rat5, session2 | 2 | 19 | 72 | 86 | 66 | 47 (32/15) | 32 | 94 | 87 |

| b | | Original | | | Rate fixed | | | Place removed | | | Rate fixed, Place randomized | | | Place removed, Rate randomized | | |
|---|---|---|---|---|---|---|---|---|---|---|---|---|---|---|---|---|
| | **Session** | **Mean** | **SE** | **P value** | **Mean** | **SE** | **P value** | **Mean** | **SE** | **P value** | **Mean** | **SE** | **P value** | **Mean** | **SE** | **P value** |
| **PRE** | **1** | -0.13 | | | 0.00 | | | -0.04 | | | -0.19 | | | -0.18 | | |
| | **2** | -0.09 | | | 0.32 | | | 0.02 | | | -0.03 | | | -0.30 | | |
| | **3** | 0.27 | | | 0.38 | | | 0.02 | | | 0.04 | | | 0.00 | | |
| | **4** | -0.32 | | | -0.51 | | | -0.34 | | | 0.31 | | | 0.03 | | |
| | **5** | 0.13 | | | -0.21 | | | 0.06 | | | -0.37 | | | 0.05 | | |
| | **6** | 0.49 | | | 0.68 | | | 0.00 | | | -0.18 | | | 0.14 | | |
| | **7** | 0.17 | | | 0.22 | | | 0.06 | | | 0.12 | | | -0.08 | | |
| | **8** | 0.02 | | | 0.27 | | | -0.05 | | | 0.06 | | | 0.23 | | |
| | **9** | -0.44 | | | -0.17 | | | -0.35 | | | 0.23 | | | -0.04 | | |
| | **10** | 0.46 | | | 0.26 | | | 0.18 | | | 0.13 | | | -0.11 | | |
| | **Mean** | 0.05 | 0.10 | 0.26 | 0.12 | 0.11 | 0.21 | -0.04 | 0.05 | 0.37 | 0.01 | 0.07 | 0.24 | -0.03 | 0.05 | 0.60 |
| **RUN** | **1** | 2.45 | | | 2.14 | | | 1.11 | | | 0.03 | | | -0.32 | | |
| | **2** | 0.74 | | | -0.19 | | | -0.11 | | | -0.34 | | | 0.05 | | |
| | **3** | 3.29 | | | 2.36 | | | 1.47 | | | -0.05 | | | -0.07 | | |
| | **4** | 2.53 | | | 2.31 | | | 1.22 | | | -0.29 | | | -0.21 | | |
| | **5** | 2.48 | | | 2.42 | | | 0.35 | | | -0.35 | | | -0.05 | | |
| | **6** | 3.15 | | | 2.29 | | | 1.46 | | | 0.09 | | | 0.05 | | |
| | **7** | 3.67 | | | 3.00 | | | 2.04 | | | 0.18 | | | 0.23 | | |
| | **8** | 2.88 | | | 2.02 | | | 1.40 | | | -0.31 | | | -0.18 | | |
| | **9** | 2.41 | | | 1.89 | | | 1.07 | | | 0.17 | | | -0.25 | | |
| | **10** | 2.53 | | | 1.80 | | | 1.67 | | | 0.10 | | | -0.14 | | |
| | **Mean** | 2.61 | 0.25 | **$9.13\times10^{-5}$** | 2.00 | 0.27 | **$1.41\times10^{-3}$** | 1.17 | 0.20 | **$1.41\times10^{-3}$** | -0.08 | 0.07 | 0.52 | -0.09 | 0.05 | 0.94 |
| **POST** | **1** | 0.72 | | | 0.35 | | | 0.26 | | | 0.17 | | | -0.09 | | |
| | **2** | 0.37 | | | 0.06 | | | -0.06 | | | -0.05 | | | -0.09 | | |
| | **3** | 1.57 | | | 1.05 | | | 0.52 | | | 0.18 | | | -0.04 | | |
| | **4** | 1.62 | | | 1.40 | | | 0.80 | | | -0.36 | | | 0.08 | | |
| | **5** | 0.70 | | | 0.78 | | | 0.18 | | | -0.13 | | | -0.07 | | |
| | **6** | 1.82 | | | 1.61 | | | 0.85 | | | -0.14 | | | -0.14 | | |
| | **7** | 2.02 | | | 1.51 | | | 1.12 | | | 0.09 | | | -0.01 | | |
| | **8** | 1.06 | | | 0.80 | | | 0.49 | | | -0.04 | | | 0.07 | | |
| | **9** | 0.68 | | | 0.42 | | | 0.21 | | | -0.05 | | | 0.10 | | |
| | **10** | 1.21 | | | 0.97 | | | 0.73 | | | -0.05 | | | 0.01 | | |
| | **Mean** | 1.18 | 0.18 | **$9.13\times10^{-5}$** | 0.90 | 0.16 | **$9.13\times10^{-5}$** | 0.51 | 0.12 | **$1.41\times10^{-3}$** | -0.04 | 0.05 | 0.94 | -0.02 | 0.03 | 0.79 |

**Table S1.**

(A) Overview of behavior, number of recorded cells and decoding accuracy for all sessions and rats.

(B) Summary statistics of the z-scored log odd difference per session and across sessions for PRE, RUN and POST.

| a | | VS Original 95% confidence interval *BOLD values statistically significant from zero* | | |
|---|---|---|---|---|
| | | **PRE** | **RUN** | **POST** |
| **Original VS Rate fixed** | **Mean AUC Diff** | 0.004 | -0.03 | -0.04 |
| | **SEM AUC Diff** | 0.03 | 0.01 | 0.01 |
| | **Lower CI** | -0.05 | **-0.04** | **-0.06** |
| | **Upper CI** | 0.05 | **-0.01** | **-0.01** |
| **Original VS Place removed** | **Mean AUC Diff** | -0.001 | -0.17 | -0.11 |
| | **SEM AUC Diff** | 0.03 | 0.01 | 0.01 |
| | **Lower CI** | -0.05 | **-0.19** | **-0.14** |
| | **Upper CI** | 0.05 | **-0.14** | **-0.09** |

| b | | VS Negative Control 95% confidence interval | | |
|---|---|---|---|---|
| | | **PRE** | **RUN** | **POST** |
| **Rate fixed VS Rate fixed + Place randomized** | **Mean AUC Diff** | 0.03 | 0.45 | 0.23 |
| | **SEM AUC Diff** | 0.03 | 0.02 | 0.01 |
| | **Lower CI** | -0.02 | **0.41** | **0.21** |
| | **Upper CI** | 0.08 | **0.48** | **0.26** |
| **Place removed VS Place removed + Rate randomized** | **Mean AUC Diff** | 0.02 | 0.31 | 0.15 |
| | **SEM AUC Diff** | 0.03 | 0.02 | 0.01 |
| | **Lower CI** | -0.03 | **0.27** | **0.12** |
| | **Upper CI** | 0.08 | **0.35** | **0.18** |

| c | DeLong Test VS Original (Significance = 0.05) | |
|---|---|---|
| | **Original VS Rate fixed** | **Original VS Place removed** |
| **PRE** | 0.88 | 0.95 |
| **RUN** | **$2.26\times10^{-6}$** | **$2.87\times10^{-52}$** |
| **POST** | **$5.04\times10^{-6}$** | **$3.10\times10^{-33}$** |

| d | DeLong Test VS Negative control (Significance = 0.05) | |
|---|---|---|
| | **Rate fixed VS Rate fixed + Place randomized** | **Place removed VS Place removed + Rate randomized** |
| **PRE** | 0.20 | 0.27 |
| **RUN** | **$1.45\times10^{-157}$** | **$5.42\times10^{-55}$** |
| **POST** | **$5.07\times10^{-69}$** | **$3.50\times10^{-31}$** |

**Table S2.**
(A) Summary statistics of the 95% confidence interval for the AUC difference between the original and the manipulated conditions.
(B) the manipulated conditions and their respective negative controls.
(C) Summary statistics of the DeLong Test comparing the ROC curves between the original and the manipulated condition.
(D) Summary statistics of the DeLong Test comparing the ROC curves between the manipulated conditions and their respective negative controls.

| | rate fixed place randomized | | | place removed rate randomized | | |
|---|---|---|---|---|---|---|
| | **Mean Difference** | **SE** | **P value** | **Mean Difference** | **SE** | **P value** |
| Sim 1 | 0.18 | 0.06 | 0.002 | 0.18 | 0.06 | 0.001 |
| Sim 2 | -0.08 | 0.12 | 0.79 | 0.01 | 0.05 | 0.79 |
| Sim 3 | -0.01 | 0.12 | 0.52 | 0.00 | 0.06 | 0.52 |
| Sim 4 | 0.05 | 0.06 | 0.79 | -0.14 | 0.07 | 0.99 |
| Sim 5 | -0.03 | 0.04 | 0.52 | -0.02 | 0.05 | 0.52 |
| Sim 6 | 0.03 | 0.09 | 0.52 | -0.08 | 0.09 | 0.71 |
| Sim 7 | -0.02 | 0.13 | 0.52 | -0.03 | 0.05 | 0.48 |
| **Sim 8** | **0.01** | **0.07** | **0.24** | **-0.03** | **0.05** | **0.60** |
| Sim 9 | 0.08 | 0.11 | 0.09 | 0.01 | 0.05 | 0.06 |
| Sim 10 | -0.03 | 0.09 | 0.55 | -0.03 | 0.05 | 0.26 |

**Table S3.**
The summary statistics for the z-scored log odd difference from 10 negative control simulations for PRE. The negative control manipulation used for main result was selected based on the median difference value out of 10 simulations.
